# Distinct denitrification phenotypes in closely related bacteria: clues to understanding variations in nitrite accumulation among *Stutzerimonas* strains

**DOI:** 10.64898/2025.12.02.691782

**Authors:** Martin Menestreau, Daniel A. Milligan, Louise B. Sennett, Linda Bergaust, Lars R. Bakken, Gary Rowley, Morten Kjos, James P. Shapleigh, Åsa Frostegård

**Affiliations:** Faculty of Chemistry, Biotechnology, and Food Sciences, Norwegian University of Life Sciences, Norwegian University of Life Sciences, Ås, Norway; School of Biological Sciences, University of East Anglia, Norwich, United Kingdom; Department of Microbiology, Cornell University, Ithaca, NY, USA

**Author notes:** Correspondance: Åsa Frostegård. **Author details:** Martin Menestreau,; Daniel A. Milligan,; Louise B. Sennett,; Linda Bergaust,; Lars R. Bakken,; Gary Rowley,; Morten Kjos,; James P. Shapleigh,; Åsa Frostegård.

**Keywords:** Denitrification phenotypes, Nitrite accumulation, *Stutzerimonas* strains

## Abstract

Nitrite (NO_2_^−^) is a key denitrification intermediate, formed from nitrate (NO_3_^−^). Transient NO_2_^−^ accumulation varies among denitrifiers, yet the underlying causes remain poorly understood, despite its potential toxicity and role in NO and N_2_O emissions. We profiled eighteen related *Stutzerimonas* strains, including the model *S. perfectomarina* ZoBell, and identified three phenotypic clusters (full, partial and low nitrite accumulators; FNA, PNA and LNA) based on the fraction of NO_3_^−^-N transiently accumulated as NO_2_^−^. LNA strains lack or express the membrane-bound nitrate reductase (NarG) late, relying on periplasmic NapA for NO_3_^−^ reduction, possibly explaining their balanced NO_2_^−^ production/reduction. FNA and PNA strains possess NapA and NarG but differ in their nitrite reductase (NirS) clades. Delayed *nirS* transcription as FNA strains transition to NO_3_^−^ respiration likely accounts for some NO_2_^−^ accumulation. However, addition of NO_3_^−^ halted NO_2_^−^ reduction in FNA strains, suggesting additional metabolic control. This may require the cytochromes NirTB, which are only found in FNA strains. The regulator DnrE was also unique to NO_2_^−^-accumulators, likely having a role in finetuning NO_2_^−^ regulation. Our findings reveal diverse NO_2_^−^-handling phenotypes among denitrifiers and provide insights for optimizing wastewater nitrogen removal and soil bioaugmentation strategies to mitigate N_2_O emissions.

## Introduction

Denitrification is one of the main pathways driving the global nitrogen cycle. Complete denitrification is the four-step reduction of nitrate (NO_3_^−^) to dinitrogen (N_2_) via the intermediate products nitrite (NO_2_^−^), nitric oxide (NO) and nitrous oxide (N_2_O). In this process, the denitrification enzymes Nar, Nir, Nor and Nos are responsible for reduction of NO_3_^−^, NO_2_^−^, NO and N_2_O, respectively (Berks et al. 1995; Zumft 1997). A wide variety of facultative anaerobic organisms, including bacteria, archaea and fungi, can perform all or some of the reduction steps associated with denitrification (Shapleigh 2013; Roothans et al. 2025). A shared trait amongst all denitrifiers is that O_2_ is the preferred electron acceptor, but if the O_2_ concentration decreases, denitrifiers use a variety of regulatory mechanisms to switch from O_2_ to N-oxides as electron acceptors (Spiro 2007; Bakken et al. 2012), leading to large variations in their transient accumulation of denitrification intermediates (Lycus et al. 2017). Current knowledge of how denitrifiers control the transition from aerobic respiration to denitrification is largely based on studies of a few model organisms capable of complete denitrification of NO_3_^−^ to N_2_. These studies have resulted in a relatively clear, though likely incomplete, understanding of transcriptional and post-transcriptional mechanisms influencing denitrification phenotypes (Spiro, 2012; Vaccaro et al. 2016; Gaimster et al. 2018).

One knowledge gap is the understanding of mechanisms controlling NO_2_^−^ accumulation in denitrifying bacteria (Liu et al. 2013). Nitrite is a central molecule in the nitrogen cycle that can be produced and consumed in many redox processes, and its reduction is the defining reaction of denitrification (Kuypers et al. 2018). In its protonated form, HNO_2_, it can diffuse into cells and cause damage itself or through the formation of toxic, reactive nitrogen species (Chislett et al. 2022). Moreover, accumulation of NO_2_^−^ in the environment can cause increased emissions of the greenhouse gas N_2_O by impairing N_2_O reduction (Highton et al. 2022) or by producing N_2_O through reactions with ferrous iron or organic matter compounds (Van Cleemput and Samater 1995). Paradoxically, in some systems, NO_2_^−^ accumulation is desirable. One example is denitratation in wastewater treatment plants, where denitrification is used instead of ammonia oxidation to provide nitrite for the anammox process (Zhuang et al. 2022).

The best-studied denitrifying bacteria, *Paracoccus denitrificans*, *Pseudomonas aeruginosa* and *Pseudomonas stutzeri* ZoBell, all transiently accumulate some NO_2_^−^ during anaerobic growth on NO_3_^−^ (Carlson and Ingraham 1983). However, studies extending beyond these model organisms have revealed substantial variations in NO_2_^−^ accumulation among diverse taxa (Lycus et al. 2017). A comparison of *Thauera* strains (Liu et al. 2013) showed that some exhibited a rapid and complete onset (RPO) of all four steps of the denitrification pathway, resulting in little to no NO_2_^−^ accumulation. Others displayed a progressive onset (PO) in which all available NO_3_^−^ was reduced to NO_2_^−^ prior to any reduction of NO_2_^−^ took place, leading to NO_2_^−^ accumulation. A shared feature of the RPO strains was that they carried the periplasmic NapA for dissimilatory NO_3_^−^ reduction but lacked the membrane-bound NarG, while the PO strains carried both NapA and NarG. All strains induced NO_3_^−^ reduction as O_2_ approached depletion, but the regulatory dynamics varied. In the RPO strains, transcription of the *nirS* and *nosZ* genes - which encode Nir and Nos, respectively – was initiated concurrently with Nar. In contrast, the NO_2_^−^ accumulating PO strains delayed transcription of these genes until all available NO_2_^−^ had been reduced to NO_3_^−^. This led to the tentative suggestion that a NO_3_^−^ responsive mechanism might regulate *nir*-transcription in the NO_2_^−^ accumulating strains, although no direct evidence for this was found (Liu et al. 2013).

The present study was initiated to clarify how regulatory mechanisms drive variations in denitrification phenotypes, and how these can differ even among closely related bacteria, with a particular focus on NO_2_^−^ accumulation. We compared eighteen strains within the former *Pseudomonas stutzeri* species cluster, now reclassified as the genus *Stutzerimonas* (Lalucat et al. 2022). This complex contains metabolically versatile organisms that are widespread in various environments (Lalucat et al. 2006). We combined genome analyses with determinations of high-resolution denitrification kinetics and transcription patterns of denitrification related genes during and after transition from aerobic to anaerobic respiration. Using deletion mutations, we investigated whether the Crp/Fnr-like regulator DnrE, known to affect transcriptional regulation in *S. stutzeri* (Härtig et al. 1999), influences NO_2_^−^ accumulation in a closely related strain. Moreover, we explored the potential role of sRNAs, which can act as both activators and repressors of transcription and translation and have been shown to influence denitrification via control of NirR in *P. denitrificans* (Gaimster et al. 2018; Moeller et al. 2021). We also considered whether electron competition between electron pathways to the different denitrification reductases (Cole 2025; Richardson 2025) could explain differences in NO_2_^−^ accumulation among the *Stutzerimonas* strains. Electrons are delivered from the TCA cycle via dehydrogenases, such as NADH or FDH dehydrogenase, to the quinone/quinol pool, which serves as a key branching point in the electron transport chain. In complete denitrifiers, electrons are then directed either to NO_2_^−^ reductases (NapA, NarG) or to the *bc*_1_ complex, a second branching point, from which electrons are passed to Nir, Nor, and NosZ via cytochromes or azurins/pseudoazurins. Competition occurs primarily at these branching points (Chen and Strous, 2013; Mania et al. 2020). In this study, we specifically investigated the electron competition between NO_3_^−^ and NO_2_^−^ reduction pathways.

Our results categorize the eighteen strains into three distinct phenotypic groups showing full, partial or low NO_2_^−^ accumulation (FNA, PNA and LNA). A major difference among these phenotypes was the presence and regulation of dissimilatory NO_3_^−^ reductases (Nar/Nap), with Nar either absent or showing delayed transcription in LNA strains. Among the NO_2_^−^-accumulating strains (FNA and PNA), differences were associated with *nir* gene cluster arrangement and NirS clade affiliation. Additional differences among phenotypic groups were observed in the transcriptional patterns of denitrification reductase genes and in the metabolic control of NO_3_^−^ and NO_2_^−^ reduction pathways. These findings indicate that denitrification phenotypes cannot be predicted solely from gene content but are instead governed by a multilayered regulatory network operating at both transcriptional and post-transcriptional levels. Since NO_2_^−^ is a central molecule in the biochemical N-cycle and is utilized by several biological processes, this knowledge is important for developing biotechnological approaches, such as using denitrifiers in combination with anammox bacteria for improved wastewater treatment technologies, or engineering the soil microbiome in agriculture to reduce greenhouse gas emissions.

## Materials and Methods

### Strains and growth conditions

Eighteen bacterial strains, previously classified as belonging to the *P. stutzeri* species complex (Lalucat et al. 2006; Li et al. 2022), were selected for comparison of their denitrification phenotypes. The strains were obtained from various culture collections. A recent taxonomic revision of the genus *Pseudomonas* reclassified these strains into several species of the novel genus *Stutzerimonas* (Lalucat et al. 2022; Gomila et al. 2022). A list of the strains, including their origin of isolation and culture collection accession numbers, is found in Table S1.

All strains were incubated at 20°C in 120 mL serum vials containing 50 mL Sistrom’s mineral medium (Sistrom 1960; Bergaust et al. 2010) supplemented with 2 mM KNO_3_ or KNO_2_, added as filter sterilized solutions. Vials prepared for cultivation under limited O_2_ conditions were sealed with gas tight butyl rubber septa and aluminum caps (Matriks AS, Norway), and air was evacuated from the 70 mL headspace by six repetitions of applying vacuum for 360 s and He overpressure for 30 s. The overpressure was released prior to the injection of the desired amount of O_2_ and N_2_O. Near-equilibrium between headspace and the liquid phase was maintained by vigorous stirring at 600 rpm using a Teflon-coated triangular magnetic bar. The stirring also prevented cell aggregation, ensuring that the cells were evenly dispersed and experienced the intended gas concentrations. At the start of each experiment, the vials were inoculated with 1 mL of aerobically growing bacterial cultures to an initial OD_600_ of 0.05-0.10. The cultures had been raised from frozen stocks and pre-cultured with vigorous stirring under fully oxic conditions for at least 4 generations to minimize the amounts of previously synthesized denitrification enzymes in the cells (Mania et al. 2020; Gao et al. 2021).

### Gas kinetics and chemical analyses

Headspace gases were monitored using a robotized incubation system as described in Molstad et al. (2007) and Molstad et al. (2016). Briefly, gas samples were taken automatically by an autosampler (Agilent GC Sampler 80) through the rubber septa of the vials using a peristaltic pump. The samples were then pumped into a gas chromatograph (Agilent 7890A) for analysis of O_2_, N_2_O, N_2_ and CO_2_ and into a chemifluorescence NO analyzer (Model 200A, Advanced Pollution Instrumentation). A thermal conductivity detector was used to measure O_2_ and N_2_, while an electron capture detector was used to measure N_2_O. To maintain a slight overpressure in the vials, each 1 mL gas sample was replaced with a slightly larger volume of He at near-atmospheric pressure. Three replicate vials were measured for each treatment unless otherwise stated (n=3).

To determine NO_2_^−^ concentrations, samples were collected from a parallel set of vials to avoid sampling losses and changes in the headspace-to-liquid ratio. The samples were collected manually at regular intervals through the rubber septum using a sterile syringe. Concentrations of NO_2_^−^ were determined immediately after each sampling, using the methodology described in Lim et al. (2018), which is based on the method by Braman and Hendric (1989) and Cox (1980). Briefly, 10 µl liquid sample was injected into a purging device containing a reducing agent (NaI) in 50% acetic acid at room temperature and the NO produced was measured with a chemifluorescence NOx analyzer (Sievers NOA 280i). Samples from three replicate vials were measured (n=3). Calculations based on cubic spline interpolation allowed estimation of values between sampling times.

### Statistical analyses

Statistical analyses were performed using the base statistics software in RStudio. In brief, one-way analysis of variance (ANOVA) tests were performed to assess whether there were significant differences in the levels of denitrification intermediates produced by the different NO_2_^−^ accumulation phenotypes and by the various overexpression mutants. When differences occurred (*p* = 0.05), mean pairwise comparisons were performed using Tukey’s test (HSD) for the post hoc analysis of the ANOVA models. Normality was tested using homoscedasticity plots and the Shapiro-Wilks test of the residuals (*p* > 0.05).

### Transcription analysis

Samples for RNA extraction (approximately 5 x 10^8^ cells per sample) were taken at intervals from the liquid phase. Cells were pelleted at 10,000 g for 7 minutes at 4°C and stabilized using RNAprotect (Bacteria Reagent, Qiagen) according to the manufacturer’s protocol. Total RNA was extracted using the RNeasy Mini Kit (Quiagen). RNA concentrations were measured using the Qubit RNA BR Assay Kit (Life Technologies). Genomic DNA was removed with Turbo DNase (Life Technologies) and genomic DNA contamination was ruled out by SYBR green qPCR amplification using primers for *nosZ* (Table S2) prior to reverse transcription. The assays were run on an Applied Biosystems StepOnePlus system, and samples with values < 5 *nosZ* copies per reaction were deemed gDNA free (Lim et al. 2016). Reverse transcription was then carried out with Maxima First Strand cDNA Synthesis Kit (Thermo Scientific), the product was diluted, and gene transcripts were quantified by digital droplet PCR (ddPCR). A ddPCR reaction mix was prepared (QX200 ddPCR EvaGreen supermix; Bio-Rad). Oil droplet suspensions were prepared using a QX200 droplet generator (Bio-Rad), and PCR reactions were performed using the temperature program: enzyme activation at 95 °C for 5 min; 40 cycles of denaturation at 95 °C for 30 s and combined annealing/extension at 60 °C for 1 min; followed by signal stabilization at 4 °C for 5 min and a final step at 90 °C for 5 min. PCR products were quantified using a QX200 droplet reader (Bio-Rad) and the data was analyzed using QuantaSoft Analysis Pro 1.0.596 (Bio-Rad). All primers used are listed in Table S2.

### Deletion of *narG* and *dnrE* in *Stutzerimonas decontaminans* strain 19SMN4

The pACRISPR/pCasPA system developed by Chen et al. (2018) for genome editing in various *Pseudomonas* species was used to construct the two deletion mutants, Δ*narG* and Δ*dnrE*, in *Stutzerimonas decontaminans* 19SMN4. pACRISPR plasmids specific to the targeted gene were constructed to contain a spacer targeting the specific gene designed using the tool CRISPOR (Concordet and Haeussler 2018) and a repair template consisting of 1000-bp flanking sequences, upstream and the downstream of the gene. The pCasPA was electroporated into *S. decontaminans* strain 19SMN4 following the protocol of Choi et al. (2006), but with electroporation parameters optimized for this strain (2500V, 200Ω, 25µF, 2 mm cuvette). Colonies containing pCasPA were selected after incubation on LB agar in the presence of 50 μg mL^−1^ tetracycline. Before the second electroporation for introduction of the gene specific plasmid pACRISPR, the strain containing the pCasPA plasmid was induced for 2 hours with 2 mg mL^−1^ L-arabinose, followed by electroporation under the same conditions as the first electroporation. Transformants were then selected on 50 µg mL^−1^ tetracycline and 150 µg mL^−1^ carbenicillin plates. The deletions were validated by PCR (for primers see Table S2) and Sanger sequencing. Colonies of successfully mutated cells were restreaked one to three times on LB agar supplemented with 10% w/v sucrose to cure the plasmids from the cells.

### *Stutzerimonas perfectomarina* ZoBell sRNA overexpression mutants

Small RNAs (sRNAs) were overexpressed in *Stutzerimonas perfectomarina* ZoBell using the arabinose-inducible vector pL2020 (Sommer et al. 2017). The pL2020 plasmid was linearized by digestion with NdeI and HindIII. sRNAs were amplified from *S. perfectomarina* ZoBell genomic DNA using overlapping PCR. The resulting PCR fragments were assembled with the digested vector using the NEBuilder HiFi DNA Assembly Master Mix (New England Biolabs) and transformed into *E. coli* DH5α competent cells. Transformants were selected on LB agar plates containing 30 µg mL^−1^ chloramphenicol. Positive clones were screened by colony PCR and verified by Sanger sequencing. Constructed plasmids were subsequently electroporated into the *S. perfectomarina* ZoBell host strain using an adapted protocol (Choi et al. 2006) with optimized electroporation parameters as described above. The growth of the sRNA overexpression mutants was assessed in LB medium supplemented with 30 µg mL^−1^ chloramphenicol to maintain plasmid selection.

### Genome sequencing and assembly

DNA was extracted with the QIAmp DNA Mini Kit (Qiagen, USA) and sent to the Norwegian Sequencing Centre (University of Oslo) for TruSeq Nano preparation and Illumina MiSeq sequencing. Reads were trimmed using Nesoni clip (https://github.com/Victorian-Bioinformatics-Consortium/nesoni) and contigs were assembled using Velvet (Zerbino and Birney 2008). Sequence annotation was carried out on the RAST server (Aziz et al. 2008). NCBI accession numbers are listed in Table S1.

### Gene sequence analyses and phylogenetic determinations

Sequences coding for housekeeping genes assigned to strains in the *P. stutzeri* species complex, as well as regulatory and structural genes encoding relevant nitrogen cycle processes, were retrieved from the NCBI databases. For protein-coding genes, the match sequences that represented uninterrupted open reading frames (ORFs) were identified, and gene synteny was compared for strains for which extensive operon data was available. The phylogenetic relationship of the strains was determined based on a multilocus sequence analysis (MLSA) of the genes encoding the small subunit RNA (16S rRNA), gyrase B (*gyrB*) and RNA polymerase (*rpoD*). The sequences were complete, except in the case of 16S rRNA genes where the first 30 residues were omitted, as they were not available for strain DSM 50238. Genes were aligned using MUSCLE v. 3.8.31 (Edgar 2004) and examined before Jukes Cantor distance estimation and maximum-likelihood tree construction, using PhyMl 3.1 (Guindon et al. 2010). Amino acid-based enzyme phylogenies were calculated using Clustal Omega (Sievers et al. 2011) and phylogenetic trees were constructed using the Whelan and Goldman model in PhyMl.

## Results

### Nitrite accumulation phenotypes and phylogeny

Analyses of the denitrification phenotypes revealed significant differences in transient NO_2_^−^ accumulation among the strains (Table 1). When incubated under denitrifying conditions with initially 100 µmol vial^−1^ of NO_3_^−^ (2 mM), the full nitrite accumulator (FNA) strains reduced ≥89 % of the provided NO_3_^−^ to NO_2_^−^ before any NO_2_^−^ reduction took place (Figure 1a and Figure S1). The partial nitrite accumulator (PNA) strains (Figure 1b and Figure S2) exhibited transient NO_2_^−^ accumulation, amounting to about half of the provided NO_3_^−^ (43.7-62.2 µmol vial^−1^ of NO_2_^−^). The low nitrite accumulator (LNA) phenotype (Figure 1c and Figure S3) showed no or little NO_2_^−^ accumulation (0.2-1.6 µmol vial^−1^), except for strains 24a75 and 4C29, which transiently accumulated 11.3 and 17.1 µmol NO_2_^−^ vial^−1^, respectively.

**Figure 1.**
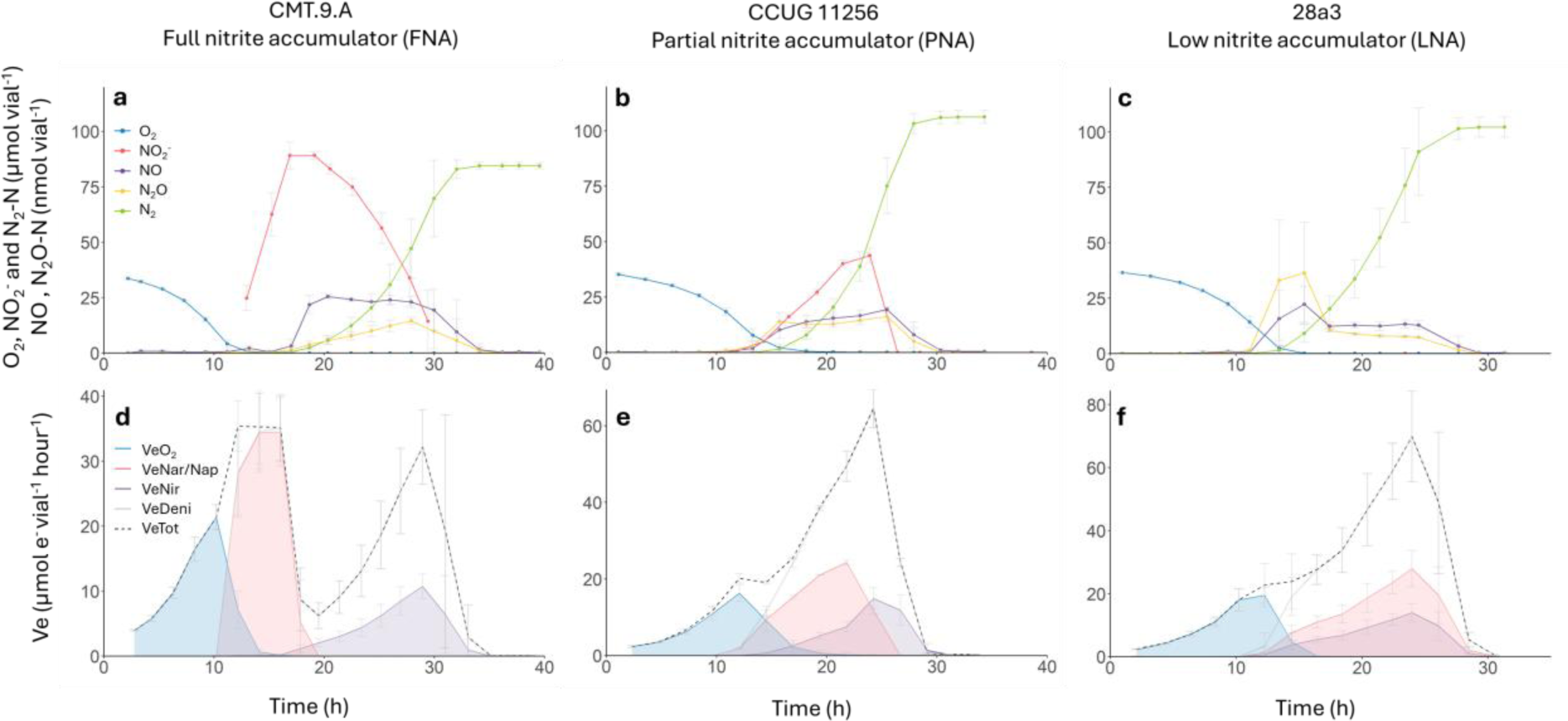
Denitrification kinetics (a–c) and electron flow rates to reductases (d–f) during and after the transition from aerobic respiration to denitrification. Strains were incubated at 20 °C in 120 mL medical vials containing 50 mL Sistrom’s mineral medium, supplemented with KNO_3_ to an initial concentration of 2 mM (100 µmol vial^−1^), with He and 1% O_2_ in the headspace. The panels show the results for three strains representing the FNA, PNA, and LNA groups. The upper panels show the measured NO_2_^−^ and gases and the lower panels show the calculated rate of electron flow to O_2_ (blue), NO_3_^−^ (red), and NO_2_^−^ (purple), with the total electron flow shown as a dashed line. All graphs show mean values from three independent replicates (n = 3); error bars represent standard deviation.

**Table 1.**
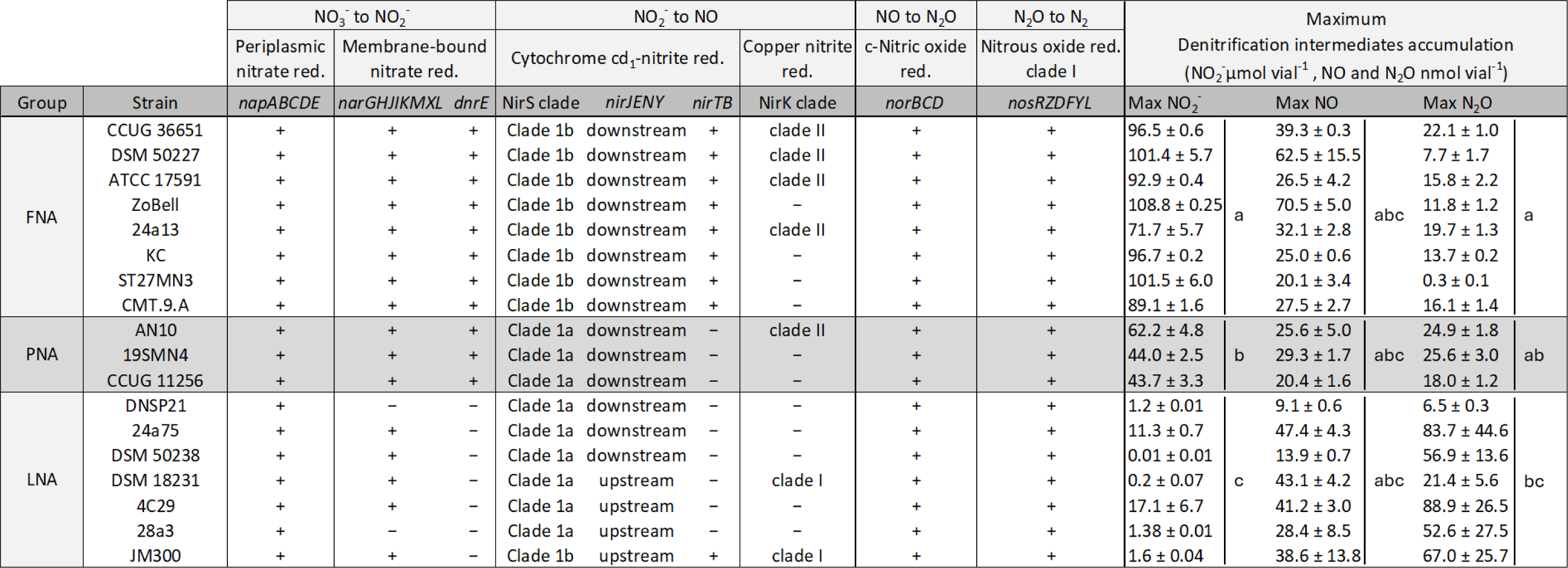
Denitrification-related genetic features and maximum accumulation of the intermediate products of 18 *Stutzerimonas* strains. The strains were characterized and grouped as full nitrite accumulators (FNA), partial nitrite accumulators (PNA) or low nitrite accumulators (LNA). The two nitrite reductases NirS and NirK were classified into different clades according to Jones et al. (2008). The genetic organization of the *nirS* related clusters *nirMCFDLGH*, located directly downstream and *nirQOP,* positioned directly upstream of *nirS*, was the same in all strains. These are therefore not mentioned in the table. The *nirJENY* cluster was found either upstream or downstream of *nirS*, depending on the strain. All *nirK* clade I genes were accompanied by the accessory gene *nirV*, located directly downstream of the *nirK* gene, while all *nirK* clade II genes were followed by an extra *norCBQD* gene cluster, not shown in the table. Statistical significance for the maximum denitrification intermediates accumulation was assessed using one-way ANOVAs (p = 0.05). Post hoc multiple comparisons of means were conducted using Tukeýs Honest Significant Difference (HSD; p = 0.05) tests. Statistical differences are indicated where the groups sharing the same letter are not statistically different.

A phylogenetic tree was constructed, incorporating the eighteen *Stutzerimonas* strains selected for analysis of denitrification phenotypes, along with additional related strains. Each straińs classification is marked in the phylogenetic tree with red (FNA), yellow (PNA) or blue (LNA) (Figure 2). The tree was based on multilocus sequence analysis (MLSA) of two complete housekeeping genes (*gyrB* and *rpoD*) and nearly complete 16S rRNA gene sequences. Only limited congruence was observed between phylogeny and the NO_2_^−^ accumulation phenotype: FNA and PNA strains clustered together and were often intertwined, whereas LNA strains were distributed across multiple, widely separated branches. The neighbor joining tree based on kmer clustering of a more extensive collection of genomes from *Stutzerimonas* strains (Figure S4) revealed a similar pattern, also consistent with other recently published phylogenetic studies (Li et al. 2022; Gomila et al. 2022).

**Figure 2.**
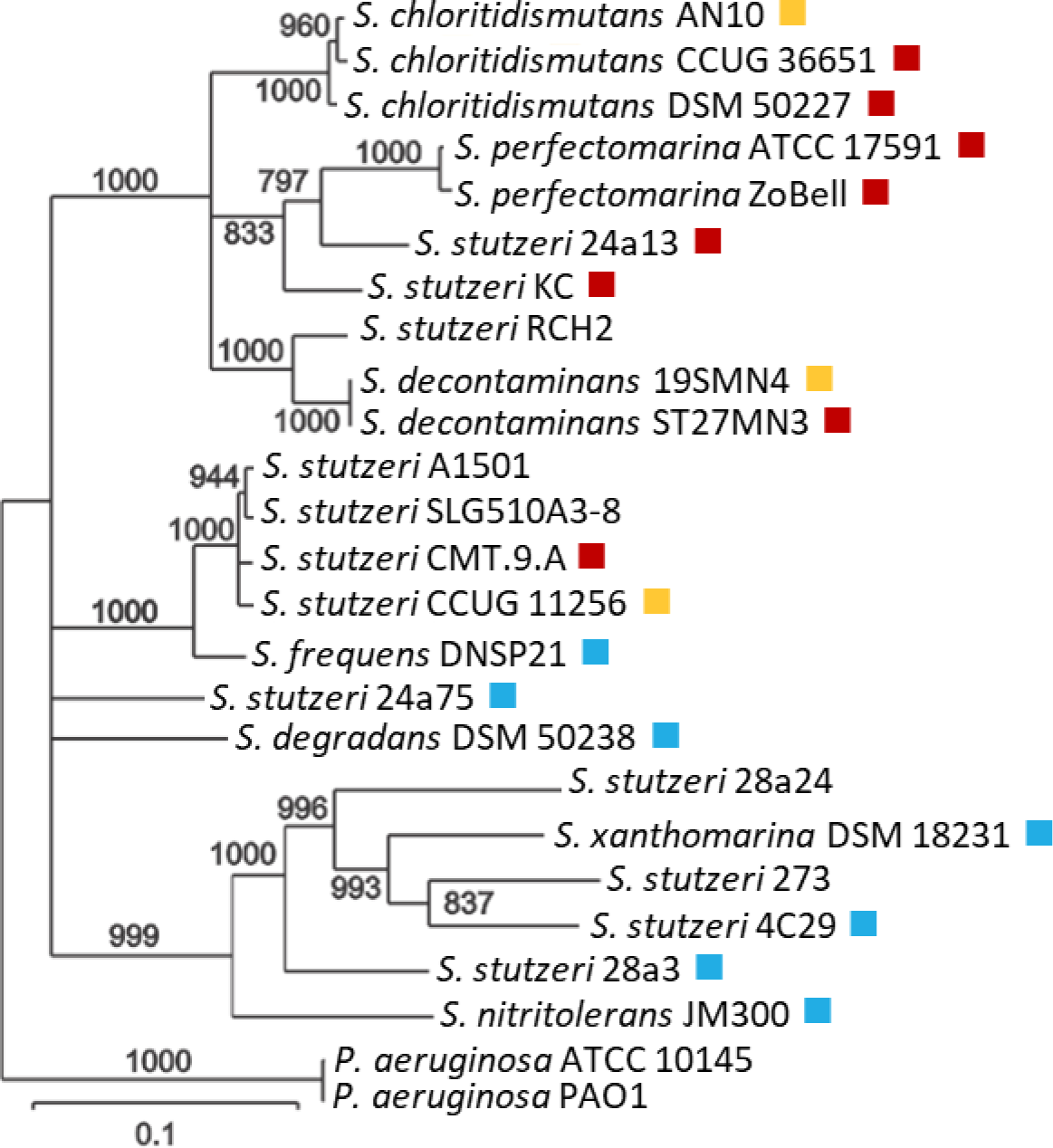
Maximum likelihood phylogenetic tree of selected *Stutzerimonas* strains and their NO_2_^−^accumulation phenotype. The three NO_2_^−^ accumulation groups are indicated by different colors in the phylogenetic tree. Red: Full NO_2_^−^ accumulators (FNA); reduced almost all available NO_3_^−^ before reducing any NO_2_^−^. Yellow: Partial NO_2_^−^ accumulators (PNA); transient accumulation of NO_2_^−^ reaching approximately 40-60% of the amount of added NO_3_^−^ Blue: Low NO_2_^−^ accumulators (LNA); 0-20% of the reduced NO_3_^−^ was transiently accumulated as NO_2_^−^. The phylogenetic analysis was a multilocus sequence analysis (MLSA) based on concatenated sequences of partial 16S rRNA and full *gyrB* and *rpoD* gene sequences. In addition to the strains analyzed for denitrification phenotypes, some other strains were included for which complete genome sequence data were publicly available. Only branches with bootstrap values ≥ 700 (of 1000) are shown. The strains were until recently included in a species complex encompassing the former *Pseudomonas stutzeri* and closely related strains (Li et al. 2022) but were recently circumscribed into several species of the novel genus *Stutzerimonas* (Laculat et al. 2022; Gomila et al. 2022). A list of the strains, including their origin and culture collection numbers, is provided in Table S1. A more detailed tree based on genome clustering, including all presently available *Stutzerimonas* genomes, is found in Figure S4.

### Selected genetic features related to denitrification

An analysis of denitrification related gene clusters was performed to identify possible relationships with the NO_2_^−^ accumulation phenotypes (Table 1). All eighteen strains had genetic potential to perform complete denitrification. They all carry the *napEDABC* gene cluster encoding periplasmic nitrate reductase (Nap). Nap also was the dominant NO_3_^−^ reductase in the clustered set of genomes (Figure S4). Two strains in the LNA group (DNSP21 and 28a3) rely only on Nap for dissimilatory NO_3_^−^ reduction, since they lack the cytoplasmic nitrate reductase Nar, encoded by the *nar* genes. The *narGHJIKMXL* cluster was found in all other strains. A notable difference in *nar*-related genes among the phenotype groups was the presence of the *dnrE* gene, which was found in all FNA and PNA strains but was absent in all LNA strains. DnrE is located within the *nar* gene cluster. It belongs to the group dissimilatory nitrate respiration regulators (DNR), a subgroup of transcriptional regulators related to the Fnr/Crp family (Trunk et al. 2010; Ebert et al. 2017), and has a potential role in regulating the *narG* gene expression (Härtig et al. 1999).

All analyzed strains carried the *nirS* gene encoding the cytochrome *cd_1_* nitrite reductase (Zumft 1997). The *nirS* gene is part of a large gene cluster encoding various enzymes involved in the synthesis and function of NirS (Jüngst et al. 1991; Philippot 2002). The FNA and PNA strains have identical *nirS* gene cluster configurations, apart from the presence of *nirTB* in the FNA strains, while the *nirS* gene cluster in the LNA strains is more diverse (Figure S5). Based on their amino acid sequences, the NirS reductases have been divided into several clades. (Jones et al. 2008; Pold et al. 2024). Following this classification, the NirS type of the FNA organisms in the present study belongs to clade 1b, while that of the PNA and LNA strains belong to clade 1a (Table 1), except the LNA strain JM300 which carries the NirS clade 1b (Figure S5). A common trait for all analyzed (Table 1) or clustered (Figure S4) *Stutzerimonas* strains with clade 1b NirS is the consistent presence of the genes *nirT* and *nirB*. NirT and NirB are tetraheme and diheme cytochromes, respectively, with a putative role in electron transfer and possibly also in the catalytic activation of the *cd_1_* cytochrome (Härtig and Zumft 1999; Zajicek et al. 2004). The *nirTB* genes are absent in all clade 1a strains.

Of the 314 genomes in the clustered tree (Figure S4), 111 carried the *nirK* gene, which encodes the copper-containing nitrite reductase NirK. Among the analyzed strains, seven possessed both *nirK* and *nirS* (Table 1). Notably, *nirK* was present in strains across all three NO_2_^−^ accumulator groups. Previously, *nirK* was divided into two main clades based on copper-binding histidine motifs: TRPHL for clade I; and SSFH(V/I/P) for clade II (Jones et al. 2008; Declayre et al. 2016). A recent study (Pold et al. 2024) proposed a more detailed classification, reassigning *Stutzerimonas* strains with clade I NirK to clade 1a, and those with clade II to clade 1e. In the analyzed *Stutzerimonas* strains carrying the *nirK* clade I gene (1a), the gene is consistently followed by *nirV*. This was observed in two of the strains in the present study, both belonging to the LNA phenotype group (Table 1). Half of the FNA strains analyzed in this study, along with one PNA strain, possess *nirK* clade II (1e), consistently followed by the *norCBQD* gene cluster.

The nitric oxide reductase cNor, which was present in all eighteen strains, is encoded by the *norCBD* operon located in the denitrification island together with the *nir* and *nos* clusters (Zumft 1997; Vollack et al. 1998). The *norCBQD* cluster present downstream of the *nirK* clade II gene in some of the FNA and PNA strains may be used under conditions when *nirK* is expressed, given their proximity. The gene cluster *nosRZDFYL* encoding the nitrous oxide reductase (NosZ) clade I was also found in the denitrification island in all the studied *Stutzerimonas* strains. The *nor* and *nos* gene cluster arrangements were identical in all strains (Table 1).

### Denitrification kinetics and transcription analysis

Electron flow rates to the different reductases were calculated from denitrification kinetics measured for all eighteen strains during and after the switch from aerobic respiration to denitrification when incubated with 1% initial O_2_ and 2 mM NO_3_^−^ (100 µmol vial⁻¹). All strains could perform the complete denitrification process, reducing NO_3_^−^ to N_2_. Examples for some representative strains are shown in Figures 1 and 3, while Figures S1, S2 and S3 show the denitrification kinetics for the remainder. Common to all strains, irrespective of which NO_2_^−^ accumulation group they belonged to, was that NO_3_^−^ reduction started when the O_2_ had decreased to 10-15 µmol vial^−1^ (corresponding to 0.6-0.9 µM in the liquid). The FNA strains reduced all the provided NO_3_^−^ to NO_2_^−^ before any NO_2_^−^ reduction took place, as shown by the lack of intermediate gas products (Figure 1a and Figure S1). The start of NO_2_^−^ reduction coincided with the peak of NO_2_^−^ accumulation and was accompanied by nearly simultaneous production of NO, N_2_O and N_2_. The sequential reduction of NO_3_^−^ and NO_2_^−^ in the FNA strains is also reflected in the electron flow kinetics (Figure 1d and Figure S1), with nearly no overlap between the curve showing the electron flow rate to NO_3_^−^ (*_eNar/Nap_V*, red shading) and that to NO_2_^−^ (*V_eNir_*; purple shading) and a clear separation between O_2_ reduction (*V_eO2_*; blue shading) and NO_2_^−^ reduction (*V_eNir_*; purple shading). The electron flow for the FNA strains typically showed a prominent dip when the cultures shifted from NO NO_3_^−^ reduction to NO_2_^−^ reduction (Figure 1d, Figure 3e and Figure S2), suggesting that only a fraction of the cells respired NO_2_^−^ as found for *Paracoccus denitrificans* (Lycus et al. 2018).

**Figure 3.**
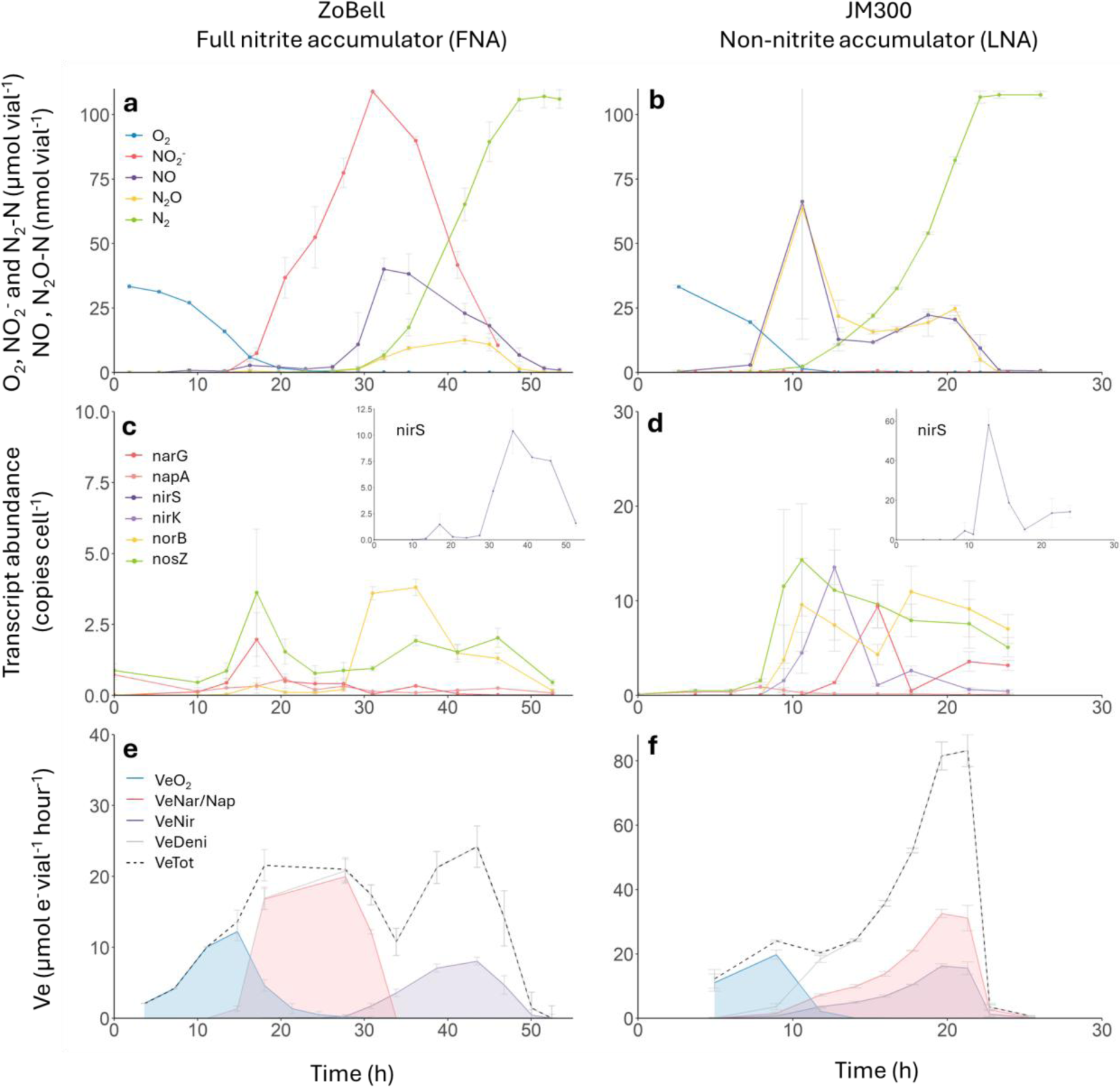
Denitrification kinetics, gene transcription and electron flow in the FNA strain ZoBell and the LNA strain JM300, switching from aerobic to anaerobic respiration. The panels show measured NO_2_^−^ and gases (a, b), the denitrification gene transcription (c, d), and electron flow rates to reductases (e, f) during and after the transition from aerobic respiration to denitrification. Strains were incubated at 20 °C in 120 mL medical vials containing 50 mL Sistrom’s mineral medium, supplemented with KNO_3_ to an initial concentration of 2 mM (100 µmol vial^−1^), with He and 1% O_2_ in the headspace. Graphs show mean values from six independent replicates (n = 6); error bars represent standard deviation.

The PNA and LNA strains reduced NO_3_^−^ and NO_2_^−^ simultaneously, as indicated by the production of NO and N_2_O, followed by the appearance of N_2_ (Figure 1b-c, Figure S2 and Figure S3). The strains CCUG 11256 (PNA) and 28a3 (LNA) were similar in their denitrification phenotypes (Figure 1b-c), except for the transient accumulation of NO_2_^−^ by the PNA strain. The simultaneous reduction of NO_3_^−^ and NO_2_^−^ was reflected in the overlap between the areas under the curves for *V_eNar/Nap_* (red shading) and *V_eNir_* (purple shading) (Figure 2e-f, Figure S2 and Figure S3). Production of N_2_ was detected somewhat later, when O_2_ was near depletion. For both the PNA and LNA groups, the electron flow indicated smooth transitions between the different electron acceptors with no distinct dips and thus an overlap between the curves for *V_eNar/Nap_* and *Ve_Nir_*.

All three NO_2_^−^ accumulation phenotypes showed little transient accumulation of NO and N_2_O (Table 1), suggesting a strong regulatory control of the production and consumption of these gaseous denitrification intermediates. When incubated with 1% initial O_2_ and 2 mM initial concentration of NO_3_^−^, the maximum net NO production was 9.1–70.5 nmol vial^−1^ with no significant (*p* = 0.598) difference between the phenotype groups in terms of maximum NO accumulation. The N_2_O accumulation remained low for all strains, ranging between 0.3 and 89 nmol vial^−1^, and was significantly lower for FNA compared to LNA (p = 0.030), while PNA exhibited intermediate levels, showing no significant difference from either FNA or LNA (p = 0.840 and 0.217).

Transcription of the denitrification reductase genes *narG, napA*, *nirS*, *norB* and *nosZ* was monitored in the strains ZoBell (FNA) and JM300 (LNA) during and after the transition from aerobic respiration to denitrification, along with the denitrification kinetics (Figure 3). In addition, transcription of the *nirK* gene, which is lacking in strain ZoBell (Table 1), was monitored in strain JM300. Strain ZoBell reduced all NO_3_^−^ (100 µmol vial^−1^) to NO_2_^−^ before any significant NO_2_^−^ reduction took place (Figure 3a), confirming its classification into the FNA group. This was also reflected in the electron flow kinetics which showed that electrons were channeled to Nir only once the electron flow to Nar stalled due to NO_3_^−^ depletion (Figure 3e). Transcription of *napA* and *nosZ* in strain ZoBell (Figure 3c) was detected already at 0 hpi (hours post inoculation). Increased transcription was observed for all the monitored genes at 10-13 hpi when less than 10 µmol of O_2_ remained in the vial, with an initial transcription peak for most genes at 18 hpi. Although transcripts for genes involved in all four denitrification steps were detected during this initial peak, it resulted in only minimal reduction activity, except for NO_3_^−^ reduction. A second pulse of transcription occurred once all NO_3_^−^ had been reduced and the NO_2_^−^ concentration reached its maximum. During this second event, the maximal levels of *norB* and *nirS* (inset) transcription were observed at 30-35 hpi. There was also a notable increase in *nosZ* transcription, although not surpassing what was measured in the initial phase. These peaks corresponded to the nearly simultaneous reduction of NO_2_^−^, NO and N_2_O (Figure 3a). Transcription was sustained until all intermediates were fully reduced to N_2_.

The LNA strain JM300 (Figure 3b, d, f) reduced NO_3_^−^ to gases, via NO_2_^−^, without any detectable net accumulation of NO_2_^−^. NO and N_2_O became detectable when the O_2_ concentration was approximately 20 µmol vial^−1^, corresponding to1.2 µM in the liquid (at 7.5 hpi), consistent with previous findings for this strain (Wittorf et al. 2018). Concentrations of these gases peaked around 11 hpi, immediately followed by N_2_ production. The simultaneous reduction of the different electron acceptors is seen in the gradual increase in total electron flow rates in JM300 as well as in the other LNA strains (Figure 3f and Figure S3). Also notable is the absence of the significant dip in the electron flow rates that was observed in most of the FNA strains during the shift from NO_3_^−^ to NO_2_^−^ reduction (Figure 3e and Figure S1). The overlapping *V* and *V_eNir_* curves (Figure 3f) reflect the simultaneous reduction of NO_3_^−^ and NO_2_^−^, also observed for the other strains characterized as LNA (Figure 2c, f and Figure S3). An important finding was that *narG* transcription was delayed by five hours compared to the other genes, indicating that NO_3_^−^ reduction was carried out by the periplasmic nitrate reductase Nap during the early phase of denitrification. A slight delay in the transcription of *nirS*, *nirK*, and *norB* was also observed compared to *napA* and *nosZ*. Like this LNA strain (JM300), delayed *narG* transcription was seen in the LNA strain DSM 50238 (Figure S6), which exhibited a transcriptional profile similar to JM300, although it lacks *nirK*.

### Effect of *narG* and *dnrE* deletions on nitrite accumulation

The wild-type PNA strain 19SMN4 carries *narG*, which is either absent (Table 1) or transcribed with a delay (Figure 3 and Figure S6) in the LNA strains, as well as *dnrE*, which is absent in all the LNA strains (Table 1). We hypothesized that deletion of either gene would shift strain 19SMN4 from the PNA to the LNA phenotype. Specifically, we predicted that NO_3_^−^ reduction rate would slow down in the *narG* mutant, as it would rely solely on the periplasmic NO_3_^−^ reductase Nap for dissimilatory NO_2_^−^ reduction, resulting in decreased NO_2_^−^ accumulation. Furthermore, given that all LNA strains lack the *dnrE* gene (Table 1), we hypothesized that the absence of this transcriptional regulator would reduce or delay the transcription of *narG* in the presence of NO_3_^−^, compared to the wild-type. This would similarly result in reduced NO_2_^−^ accumulation producing a phenotype resembling that of the LNA strains. To test these hypotheses, we generated deletion *narG* and *dnrE* mutants in the PNA strain 19SMN4.

A comparison of the denitrification phenotypes of the wild-type and the two mutants showed that the mutants remained complete denitrifiers, as all provided NO_3_^−^ was fully reduced to N_2_ (Figure 4). Both the wild-type and the mutants transitioned to denitrification, indicated by the accumulation of low amounts of NO (16.5-43.2 nmol vial^−1^) and N_2_O (11.0-34.9 nmol vial^−1^), when O_2_ fell below 7.5 µmol vial^−1^(0.45 µM in the liquid). The two mutants and the wild-type strain showed, however, different denitrification phenotypes regarding their NO_2_^−^ accumulation and the time it took to complete denitrification (seen as an N_2_ plateau). The wild-type strain accumulated approximately 50% of the provided 100 µmol NO_3_^−^ vial^−1^ as NO_3_^−^ (62.8 µmol vial^−1^), supporting its classification as PNA, determined in a separate experiment (Figure S2) where this strain accumulated 44.0 µmol vial^−1^ NO_2_^−^. In contrast to the wild-type, the Δ*narG* mutant accumulated <1.3 µmol vial^−1^ NO_2_^−^ (Figure 4). It had thus obtained an LNA phenotype similar to the strains *Stutzerimonas frequens* DNSP21 and *S. stutzeri* 28a3 (Figure 1c and Figure S3a), both of which lack *narG* and thus use only Nap for dissimilatory NO_3_^−^ reduction (Table 1). As expected, deleting *narG* slowed denitrification so that the reduction of the provided NO_2_^−^ to N_2_ took almost three times longer than for the wild-type (78 h *vs* 28 h). Knocking out the *dnrE* gene had a less severe effect on the time it took to complete denitrification (38 h). However, and contrary to our expectations, the Δ*dnrE* mutant showed increased accumulation of NO_2_^−^ compared to the wild-type, reaching 80.1 ± 4.99 µmol vial^−1^. The NO_2_^−^ accumulation rates were however almost equal with 5.5 ±0.7 µmol vial^−1^ h^−1^ for the wild-type and 5.5 ± 0.4 µmol vial^−1^ h^−1^ for the Δ*dnrE* mutant.

**Figure 4.**
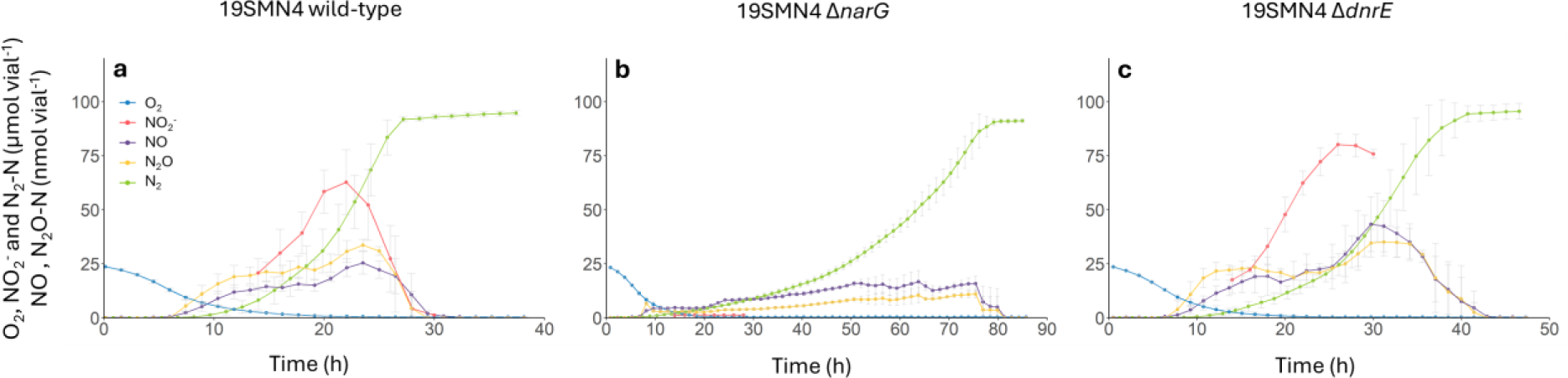
Effect of deleting *narG* and *dnrE* on the phenotype of the PNA strain *Stutzerimonas decontaminans* 19SMN4. The panels show the measured NO_2_^−^ and gases during and after the transition from aerobic respiration to denitrification for the wild type (a), *narG* deletion mutant (b) and *dnrE* deletion mutant (c). Strains were incubated at 20 °C in 120 mL medical vials containing 50 mL Sistrom’s mineral medium, supplemented with KNO_3_ to an initial concentration of 2 mM (100 µmol vial^−1^), with He and 1% O_2_ in the headspace. Graphs show mean values from three independent replicates (n = 3); error bars represent standard deviation.

### Competition for electrons between the NO_3_^−^ and NO_2_^−^ reduction pathways

To investigate whether competition for electrons between the NO_3_^−^ and NO_2_^−^ reduction pathways influences NO_2_^−^ accumulation, we added NO_3_^−^ to actively NO_2_^−^ reducing cultures. Three strains were studied (ZoBell, FNA; 19SMNA, PNA and JM300, LNA), each representing one of the phenotype groups (Figure 5a-c). The cultures were initially incubated under the denitrifying conditions described above to induce the development of the denitrification proteome, after which the cells were transferred to new anoxic vials (initial OD_600_ 0.3) with medium containing 100 µmol NO_2_^−^ vial^−1^ (corresponding to 2 mM NO_2_^−^ in the medium) (t=0). When approximately one third of the initial NO_2_^−^ had been reduced, 100 µmol vial^−1^ of NO_3_^−^ (corresponding to 2 mM NO_3_^−^ in the medium) was added to the vial. For strain ZoBell (Figure 5a), the addition of NO_3_^−^ to the actively NO_2_^−^ respiring culture led to an immediate shift to NO_3_^−^ respiration and a complete halt of the NO_2_^−^ reduction until all the added NO_3_^−^ had been reduced. This shift is also seen from the plummeting of the NO concentration and the plateauing of N_2_. Once all the NO_3_^−^ had been reduced, NO_2_^−^ respiration resumed along with the downstream denitrification steps as seen from the decrease of NO_2_^−^ and the concomitant accumulation of NO (<50 nmol vial^−1^). N_2_ increased until it reached a plateau at approximately 200 µmol N_2_-N vial^−1^, reflecting the complete reduction of the added NO_3_^−^ and NO_2_^−^.

**Figure 5.**
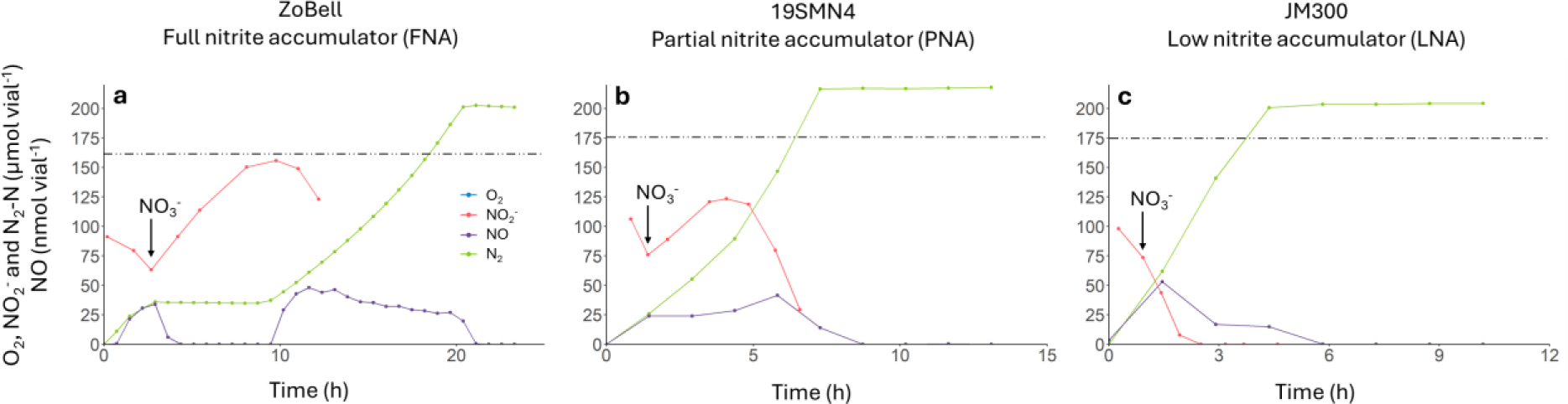
Competition for electrons between NO_3_^−^ and NO_2_^−^ reduction pathways. Results for three *Stutzerimonas* strains, each representing one of the three nitrite accumulation phenotypes (a–c). Experiments were performed in biological triplicates (n = 3). The timing of NO_3_^−^ addition varied between replicate vials; therefore, one representative replicate is shown here, while the other replicates, which displayed very similar patterns, are presented in Fig. S7. Cultures were pre-grown anoxically (100% He headspace) in Sistrom’s medium with NO_3_^−^ to induce full denitrification proteomes, then transferred at t = 0 to fresh anoxic vials containing only 2 mM NO_2_^−^ (100 µmol vial^−1^). When approximately one-third of the NO_2_^−^ was reduced, 100 µmol of NO_3_^−^ (corresponding to 2 mM in the medium) was added (indicated by the arrow).

In a similar experiment, strain 19SMN4 (PNA) reduced NO_3_^−^ and NO_2_^−^ simultaneously following supplementation with NO_3_^−^, and continuously reduced NO_2_^−^ further to N_2_, as seen by the constant increase of N_2_ and lack of an intermittent N_2_ plateau (Figure 5b). NO_3_^−^ was apparently reduced faster than NO_2_^−^, resulting in a transient accumulation of NO_2_^−^ from 75 to 123 µmol vial^−1^(thus about half of the added NO_2_^−^) until all the available NO_3_^−^ was depleted. The LNA strain JM300 (Figure 5c) continued to reduce NO_3_^−^ despite the addition of NO_3_^−^, with no accumulation of NO_2_^−^. The NO_2_^−^ reduction proceeded without any apparent reduction in rate until all NO_3_^−^and NO_2_^−^ had been consumed, seen as N_2_-N reaching a plateau. As observed with strain ZoBell, the accumulation of NO peaked at ≤ 50 nmol vial^−1^ in experiments with strains 19SMN4 or JM300 (Figure 5b-c).

### Analysis of a possible role of sRNAs in regulating nitrite accumulation

We investigated potential regulatory mechanisms underlying the nitrite accumulation phenotype by examining the role of small RNAs (sRNAs). Inclusion of selected sRNAs was based on their previous association to either regulation of the broader nitrogen cycle, to denitrification specifically, to anaerobic regulators, and to regulators located within denitrification gene clusters in *Stutzerimonas* sp. (Lin et al. 2015; Zhan et al. 2016; Wang et al. 2022). With these criteria, eight putative sRNAs (crcZ, nfiS, P11, P15, pda200, phrS, prrF1, rsmY) were selected for investigation and overexpression strains constructed. Overexpression has been used to assign function to sRNAs from a wide range of bacterial species and associated processes (Massé and Gottesman 2002; Gaimster et al. 2019). We assessed the NO_2_^−^ accumulation under anaerobic conditions for the sRNA-overexpressing strains (Figure S8). During anaerobic growth, NO_2_^−^ accumulation peaked between 83.2 ± 7.7 and 104.2 ± 0.4 µmol vial^−1^. However, pairwise Tukey HSD tests revealed no statistically significant differences in NO_2_^−^ accumulation between any of the overexpression strains and the control (p > 0.05).

## Discussion

Nitrite accumulation can have significant environmental consequences, as it plays a central role in the emissions of N_2_O, NO and HONO from various environments (Bhattarai et al. 2018; Frostegård et al. 2022; Song et al. 2023), and can also hamper N removal efficiency in wastewater plants (Zhou et al. 2011). Despite its importance, variation in patterns of transient NO_2_^−^ accumulation among denitrifying organisms has received relatively little attention, and mechanisms underlying this phenotype remain poorly understood. Our study of eighteen closely related strains revealed a greater diversity in denitrification phenotypes within the genus *Stutzerimonas* than previously recognized, with strains exhibiting full, partial, or low NO_2_^−^-accumulating phenotypes. We identified multiple factors that either individually or in concert contribute to varying levels of transient NO_2_^−^ accumulation in *Stutzerimonas*, including genetic configurations as well as regulatory mechanisms operating at both the transcriptional and post-transcriptional/metabolic levels. The phenotype grouping showed partial congruence with phylogeny (Figure 2): FNA and PNA were phylogenetically intertwined, and in some cases, even strains of the same species belonged to different phenotype groups. The LNA strains were mostly phylogenetically separated from the two other phenotype groups, with one exception (DNSP21). Genome clustering supported these observations, with the exception that strains DNSP21 and 24a75 showed proximity (Figure S4). The limited congruence between phylogeny or genome clustering and NO_2_^−^ accumulation phenotype is consistent with findings from the genus *Thauera* (Liu et al. 2013) and underscores the need for caution against inferring regulatory phenotypes based solely on taxonomic relatedness.

The genome survey of denitrification-related genes (Table 1) did not conclusively explain the three phenotype groups, as variations in the gene clusters encoding both dissimilatory NO_3_^−^ and NO_2_^−^ reduction were not consistently associated with specific phenotypes. In contrast, the clusters for NO and N_2_O reduction were arranged consistently across all three groups. A previous investigation of bacteria in the genus *Thauera* (Liu et al. 2013) suggested that *nar* plays a key role in NO_2_^−^ accumulation, since strains carrying both the *nar* and *nap* clusters accumulated NO_2_^−^, while those with only the *nap* cluster did not. In line with this, all *Stutzerimonas* strains classified as FNA and PNA in the present study carried the *nar* genes. However, unlike the *Thauera* strains, the *nar* cluster was also found in all but two of the LNA strains. This suggests that additional mechanisms controlling the expression of denitrification related genes are involved in determining the NO_2_^−^ accumulation levels.

To investigate whether transcriptional control mechanisms could explain why the LNA strains possessing both *nar* and *nap* accumulated little or no NO_2_^−^, we measured transcriptional activity in two such strains. Strain JM300 showed significantly delayed *narG* transcription by approximately 4-6 h compared to other denitrification related genes (Figure 3d), implying that it relied only on Nap for NO_3_^−^ reduction during this period. This slower NO_3_^−^ reduction allowed Nir to maintain pace, preventing NO_2_^−^ accumulation. Interestingly, strain JM300 carries the NO_2_^−^ reductase gene *nirK* in addition to *nirS*, and it cannot be excluded that this contributed to the lack of NO_2_^−^ accumulation. To explore this further, we performed a transcription analysis of the LNA strain DSM 50238, which carries both *napA* and *narG* but only *nirS* for dissimilatory NO_2_^−^ reduction (thus lacking *nirK*). These strains showed the same temporal pattern of transcription as JM300, with a significant delay in *narG* transcription (Figure S5), although copy numbers were lower for DSM 50238, for which real-time PCR was used, compared to the ddPCR analysis used for JM300. Collectively, these results reinforce the role of NarG as a primary factor in determining whether a denitrifier accumulates NO_2_^−^. This is further supported by the results from the *narG* knock-out mutant, which shifts PNA strain 19SMN4 to a LNA phenotype (Figure 4).

While the presence and activity of Nar appear to be significant factors in determining whether a strain accumulates NO_2_^−^, they do not conclusively explain the difference between the phenotype groups. A primary distinction between the FNA group and the PNA/LNA groups appears to lie in their transcriptional control of *nirS* (Figure 3). The FNA phenotype was characterized by a progressive onset of denitrification, whereas the LNA phenotype showed a rapid and complete onset, similar to reports on *Thauera* strains (Liu et al. 2013). In that study, the FNA strain *Thauera terpenica* showed a low transcription peak of *nirS* shortly before O_2_ depletion, but the main transcription was delayed and coincided with the peak of NO_2_^−^ accumulation and depletion of NO_3_^−^. This precise timing of *nirS* transcription led to the tentative suggestion that NO_3_^−^ interacted with one of the regulatory proteins involved in *nirS* transcription, possibly a repressor protein, though no evidence was provided to support such a mechanism. The present, more detailed analysis revealed a similar transcription pattern in *S. perfectomarina* strain ZoBell. Taken together, the findings suggest that temporal control of transcription plays a major role in the near complete NO_2_^−^ accumulation in FNA strains, although other control mechanisms are likely at play as well.

The FNA strains showed an almost complete inhibition of NO_2_^−^ reduction while NO_3_^−^ was present, whereas the PNA strains reduced NO_2_^−^ in the presence of NO_3_^−^, albeit not at a rate sufficient to immediately reduce the produced NO_2_^−^ as effectively as the LNA strains (Figure 5a-c). This suggests that, in both PNA and LNA organisms, electron flow can occur simultaneously to Nar/Nap and Nir, enabling some Nir activity even when NO_3_^−^ is available. The observed difference in NO_2_^−^ accumulation between FNA and PNA strains is unlikely to be explained by the electron pathway to NarG, via NarHI, being a stronger competitor for electrons than the *bc_1_* complex in FNA organisms relative to PNA organisms. One notable difference between these two groups is the clade of NirS, with FNA strains possessing clade 1b while PNA strains have clade a. This distinction is accompanied by the presence of the genes *nirT* and *nirB* in the clade 1b cluster, but not in the clade 1a cluster (Table 1). These genes encode tetraheme and diheme *c*-type cytochromes, respectively. NirT is a member of the NapC/NirT family of cytochromes, which are characterized by their electron transfer capacity. In addition, NirT may catalytically activate cytochrome *cd₁*, potentially serving as an essential activating partner for NirS (Zajicek et al. 2004). Notably, NirT is needed for functional NO_2_^−^ reduction in *S. perfectomarina* strain ZoBell (Jüngst et al. 1991; Härtig and Zumft 1999). The NirT and NirB proteins can apparently form a functional heterodimer, as fused NirTB has been found in other NO_2_^−^ accumulating bacteria, such as *Halomonas* sp. 2A, which was detected in a denitritation bioreactor where denitrifying bacteria supplied NO_2_^−^ for anaerobic ammonium oxidation (Li et al. 2018). The involvement of NirTB in NO_2_^−^ reduction is supported by the location of their genes immediately downstream of *nirS*. However, despite these indications of function, the extent to which NirT and NirB contribute to the near-complete inhibition of NirS activity by NO_3_^−^ in FNA strains remains unresolved.

Another difference between the phenotype groups was that the *dnrE* gene was present in all FNA and PNA strains but absent in all LNA strains (Table 1), making it a candidate gene for involvement in NO_2_^−^ accumulation. DnrE belongs to the FNR/CRP family of transcriptional regulators, which sense environmental signals such as O_2_ and NO. We hypothesized that knocking out *dnrE* in an FNA or PNA strain would decrease or delay the transcription of *narG*, thereby slowing NO_3_^−^ reduction and reducing NO_2_^−^ accumulation. Contrary to our expectations, the NO_2_^−^ accumulation rate of the PNA strain 19SMN4 deletion mutant remained equal to the wild-type (Figure 4), excluding DnrE from playing a direct regulatory role in NO_2_^−^ accumulation. The *dnrE* gene is located within the *nar* gene cluster, alongside *narXL*. NarXL is a two-component sensor–regulator system that senses NO_3_^−^ and NO_2_^−^ with a higher affinity for NO_3_^−^, which directly controls the expression of *narG* and *dnrE* (Härtig et al. 1999). Consequently, in the *dnrE* knockout mutant, *narG* expression was maintained, possibly through NarXL, despite the absence of *dnrE*, which makes it challenging to elucidate the specific role of *dnrE* in regulating NO_2_^−^ accumulation. Despite high homology among the Dnr regulatory proteins DnrE, DnrS and DnrD (>40% identity and >90% coverage), their functions are distinct: DnrS senses O_2_, DnrD senses NO, while the role of DnrE remains unknown. The regulation of *dnrE* appears to be conserved in all FNA and PNA strains, as indicated by a highly conserved upstream intergenic region containing two putative NarL binding sites, a ribosome binding site (RBS), and two promoters (Figure S9). This conservation suggests a similar regulatory mechanism controlling *dnrE* expression. Consistent with Vollack et al. (1999), no FNR-box was detected upstream of *dnrE*, suggesting that the main anaerobic regulator, Anr, which senses O_2_, may not directly regulate *dnrE*. Nevertheless, *dnrE* transcription increases in the presence of both NO_3_^−^ and O_2_, indicating that another oxygen-sensitive factor may trigger its expression. This suggests that DnrE is involved in a complex, multilayered regulatory network (Spiro 2012).

In our search for additional regulatory mechanisms influencing denitrification, we examined the potential role of small RNAs (sRNAs) in modulating NO_2_^−^ accumulation in the FNA strain *S. perfectomarina* ZoBell. Candidate sRNAs were selected based on their conservation across *Pseudomonas* species, their genomic proximity to denitrification gene clusters, and previous associations with nitrogen metabolism. Although several candidates were successfully overexpressed under anaerobic conditions, no significant changes in NO_2_^−^ accumulation were observed, suggesting that none of the selected sRNAs directly regulates denitrification under the tested conditions although mutational analyses would be needed to fully rule them out. These findings highlight the limitations of a candidate-based approach and emphasize the need for broader transcriptomic analyses. In particular, *de novo* RNA-seq analyses, comparing closely related *Stutzerimona*s strains exhibiting different NO_2_^−^ accumulation phenotypes, could provide valuable insight into the denitrification specific transcriptional landscape.

Taken together, the results enhance the understanding of how variations in regulatory phenotypes may influence NO_2_^−^ accumulation across various environments and highlight how organisms optimize energy savings by temporarily blocking steps in the denitrification pathway, albeit at the cost of providing substrates to competing populations. Importantly, these findings have significant implications for various biotechnological approaches. FNA strains can supply NO_2_^−^ for denitratation in anammox wastewater treatment (Zhuang et al. 2022). NNA strains, on the other hand, may be candidates for engineering soil microbiomes for N_2_O mitigation (Hiis et al. 2024), since they maintain low levels of all denitrification intermediates.

## Conclusions

Our study demonstrates substantial variation in denitrification phenotypes among closely related *Stutzerimonas* strains, driven by a complex interplay of genetic, transcriptional, and post-transcriptional mechanisms that influence NO_2_^−^ accumulation. We found that even organisms with a complete set of denitrification genes can act as partial denitrifiers by temporarily constraining specific steps, as seen in the FNA strains. This likely conserves energy, providing an advantage in environments with fluctuating O_2_ levels where full denitrification is only occasionally required. Similar to classical “partial denitrifiers”, which lack certain genes and therefore share metabolic tasks (Roothans et al. 2025), FNA-like organisms rely on community members capable of consuming NO_2_^−^, including other denitrifiers, DNRA organisms (performing dissimilatory NO_3_^−^/NO_2_^−^ reduction to ammonia), or NO_2_^−^ oxidizers if O_2_ is available, to keep NO_2_^−^ concentrations non-toxic. This collaboration helps prevent the buildup of NO_2_^−^ and the formation of environmentally harmful byproducts such as NO and N_2_O, including those from abiotic NO_2_^−^ reactions (Lim et al. 2018). While the presence and timing of *narG* transcription were identified as key determinants of transient NO_2_^−^ accumulation, other factors—including the type of *nirS* and associated *nirT/nirB* genes, as well as competition for electrons between Nar and Nir pathways—also play important roles. Our results highlight the limitations of predicting NO_2_^−^ accumulation potential based solely on taxonomy or gene presence. Overall, these findings underscore the complexity of denitrification regulation and emphasize the need for integrative approaches combining microbial physiology, biochemistry, and transcriptomics to link genotype to denitrification phenotypes. A deeper understanding of these mechanisms is essential for managing NO_2_^−^ and related intermediates in both environmental and engineered systems, such as wastewater treatment, and for supporting efforts to mitigate greenhouse gas emissions.

## Author contributions

**Martin Menestreau:** investigations, formal analysis, visualization, data interpretation, writing – original draft, writing – review and editing. **Daniel A. Milligan:** preliminary investigations (gas kinetics, transcription, genome analyses), writing – early draft. **Louise B. Sennett:** data analysis (statistics), writing-- editing. **Linda Bergaust:** data interpretation, writing – review and editing. **Lars R. Bakken** data interpretation, writing – review and editing. **Gary Rowley:** supervision of sRNA work, writing – review and editing. **Morten Kjos:** supervision of CRISPR analyses, writing – review and editing. **James P. Shapleigh:** data analysis (phylogeny), data interpretation, validation, writing – review and editing. **Åsa Frostegård:** conceptualization, funding acquisition, supervision, project administration, data interpretation, validation, writing – original draft, writing – review and editing. All authors contributed to the article and approved the submitted version.

## Acknowledgements

This work was supported by the Research Council of Norway, project no. 325770 (awarded to Å.F.).

## Data availability statement

The gas kinetics and transcription data that support the findings of this study are available on request from the corresponding author.

