## Supplementary Figures S1, S2, S3, S5, S6, S7, S8, S9; Tables S1, S2, S3; Fig S4S for "Distinct denitrification phenotypes in closely related bacteria: clues to understanding variations in nitrite accumulation among *Stutzerimonas* strains"

Figure S1. Denitrification kinetics and electron flow rates in FNA strains during and after the transition from aerobic respiration to denitrification

Figure S2. Denitrification kinetics and electron flow rates in PNA strains during and after the transition from aerobic respiration to denitrification

Figure S3. Denitrification kinetics and electron flow rates in NNA strains during and after the transition from aerobic respiration to denitrification

Figure S4. See separate file.

Figure S5. Maximum likelihood phylogeny of full-length NirS amino acid sequences in *Stutzerimonas* strains and gene arrangement of the *nirS* gene clusters

Figure S6. Denitrification kinetics (a), denitrification gene transcription (b), and electron flow rates to reductases (c) during and after the transition from aerobic respiration to denitrification for the NNA strain DSM 50238

Figure S7. Competition for electrons between $NO_3^-$ and $NO_2^-$ reduction pathways in the three Stutzerimonas strains, each representing the three groups of nitrite accumulators

Figure S8. Anaerobic nitrite accumulation kinetics of *Stutzerimonas perfectomarina* ZoBell sRNA overexpression mutants

Figure S9. MAFFT multiple sequence alignment of the intergenic region upstream of *dnrE* in 11 Stutzerimonas strains

### Supplementary Tables

Table S1. Description of the *Stutzerimonas* strains used in this study.

Table S2. Primers used in this study.

Table S3. Bacterial strains and plasmids used for generating *Stutzerimonas decontaminans* deletion mutant and the *Stutzerimonas perfectomarina* overexpression mutants in this study.

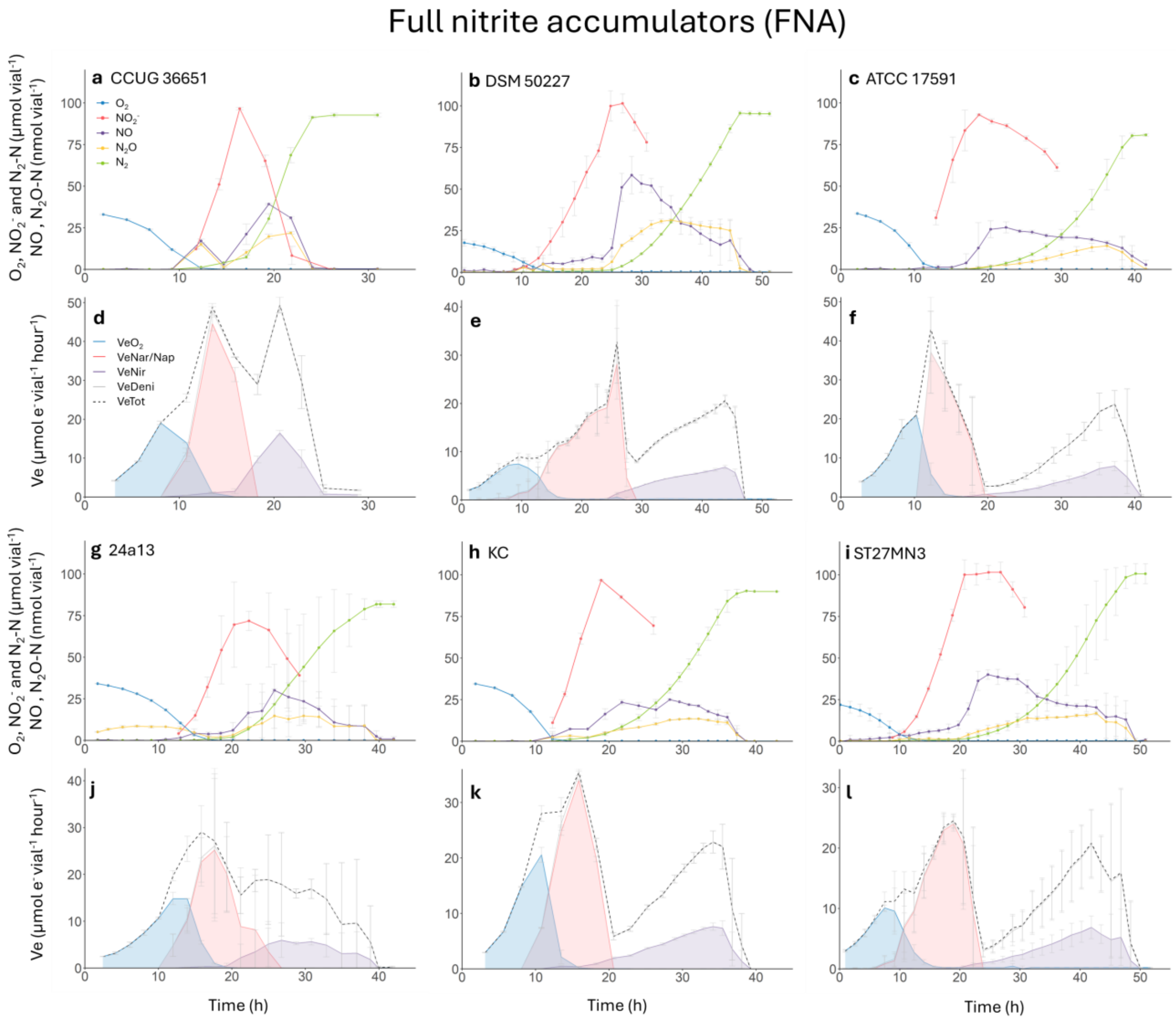


**Figure S1. Denitrification kinetics and electron flow rates in FNA strains during and after the transition from aerobic respiration to denitrification**. Results for six *Stutzerimonas* strains representing the FNA (full nitrite accumulator) group. The strains were incubated at 20 °C in 120 mL medical vials containing 50 mL Sistrom's mineral medium supplemented with $KNO_3^-$ to an initial concentration of 2 mM (100 µmol vial$^{-1}$), with He and 1% $O_2$ in the headspace. For each strain, the upper panel shows the measured amounts of each gas, and the lower pane shows the calculated rates of electron flow to $O_2$ (blue), to $NO_3^-$ (red), and to $NO_2^-$ (purple). The dashed lines show the total electron flow. All graphs show mean values from three independent replicates (n = 3); error bars indicate standard deviation.

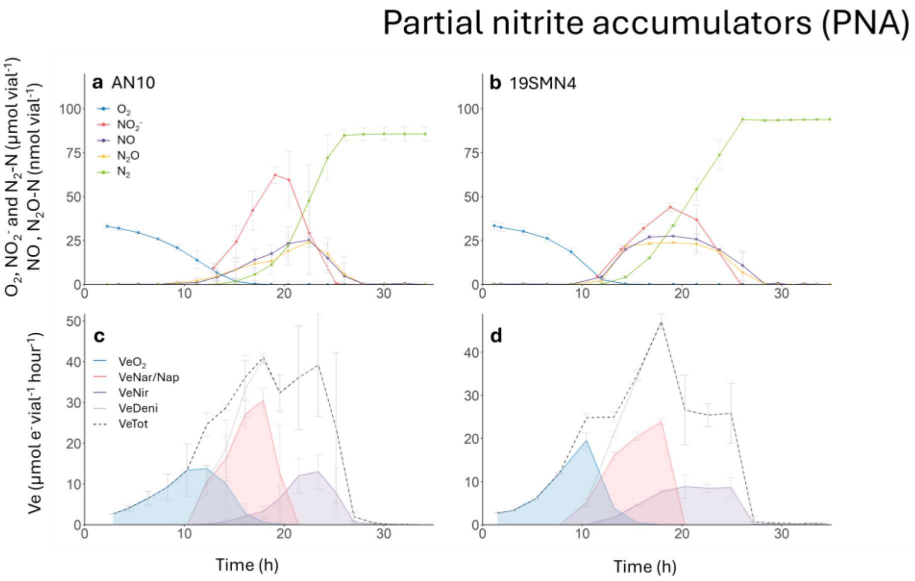


**Figure S2. Denitrification kinetics and electron flow rates in PNA strains during and after the transition from aerobic respiration to denitrification** Results for two *Stutzerimonas* strains representing the PNA (partial nitrite accumulator) group. The strains were incubated at 20 °C in 120 mL medical vials containing 50 mL Sistrom's mineral medium supplemented with $KNO_3^-$ to an initial concentration of 2 mM (100 µmol vial$^{-1}$), with He and 1% $O_2$ in the headspace. The shaded areas in panels c–d represent electron flow to $O_2$ (blue), $NO_3^-$ (red), and $NO_2^-$ (purple). All graphs show mean values from three independent replicates (n = 3); error bars indicate standard deviation. The denitrification kinetics of *Stutzerimonas decontaminans* 19SMN4 were characterized in two independent experiments (see Fig. 4), both of which exhibited identical phenotypes.

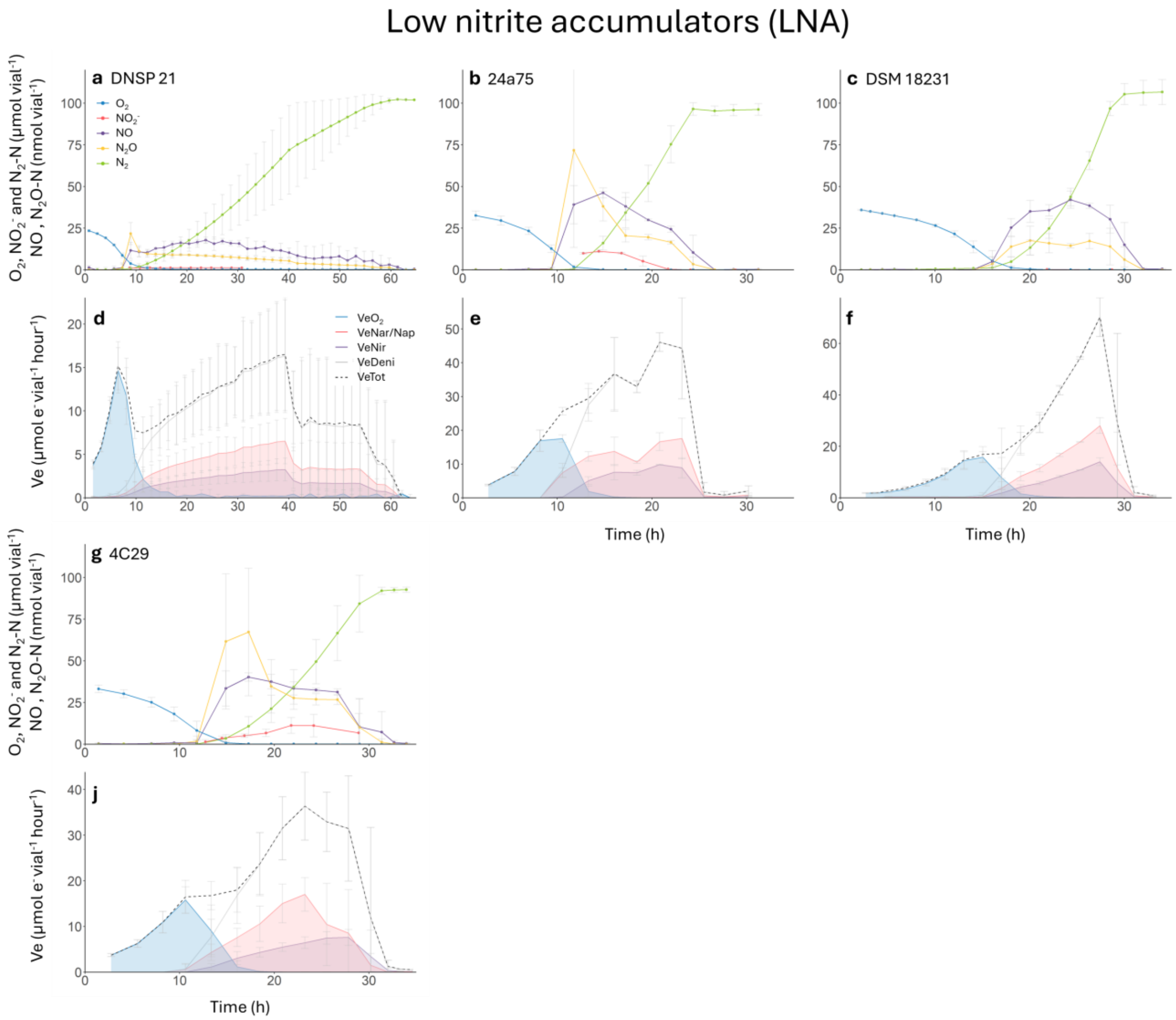


**Figure S3. Denitrification kinetics and electron flow rates in LNA strains during and after the transition from aerobic respiration to denitrification.** Results for five *Stutzerimonas* strains representing the LNA (low nitrite accumulator) group. The strains were incubated at 20 °C in 120 mL medical vials containing 50 mL Sistrom's mineral medium supplemented with $KNO_3^-$ to an initial concentration of 2 mM (100 μmol vial⁻¹), with He and 1% $O_2$ in the headspace. The shaded areas in panels d–f and j represent electron flow to $O_2$ (blue), $NO_3^-$ (red), and $NO_2^-$ (purple). All graphs show mean values from three independent replicates (n = 3); error bars indicate standard deviation.

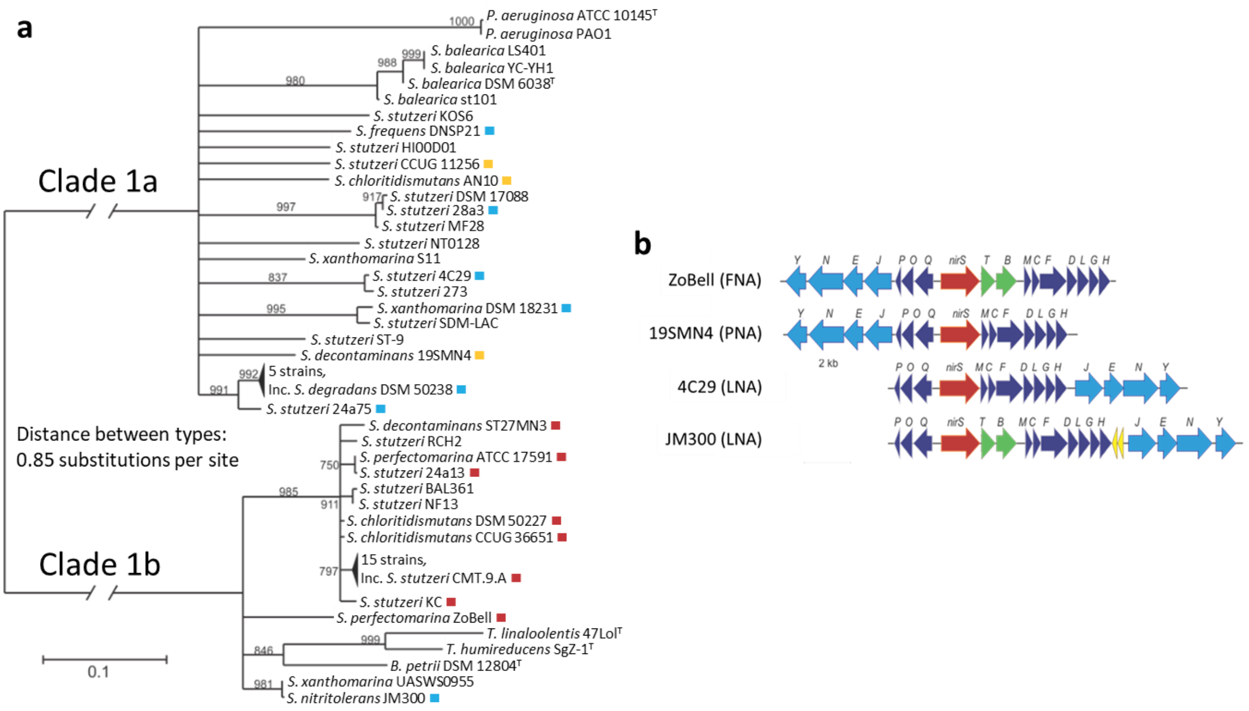


**Figure S5. Maximum likelihood phylogeny of full-length NirS amino acid sequences in *Stutzerimonas* strains and gene arrangement of the *nirS* gene clusters.** Only branches with bootstrap values ≥ 700 (of 1000) are shown. Strains are separated into two NirS types, termed clade 1a and clade 1b, according to the classification by Pold et al. (2024). Red, yellow, and blue indicate the three $NO_2^-$ accumulation phenotypes: FNA, PNA, and LNA, respectively. Panel b shows representative arrangements of the *nirS* gene cluster identified among the 18 studied *Stutzerimonas* strains, illustrated using four strains corresponding to the three $NO_2^-$ accumulation phenotypes: FNA (ZoBell), PNA (19SMN4), and LNA (4C29 and JM300). Uncharacterized ORFs with non-conserved synteny are shown in yellow.

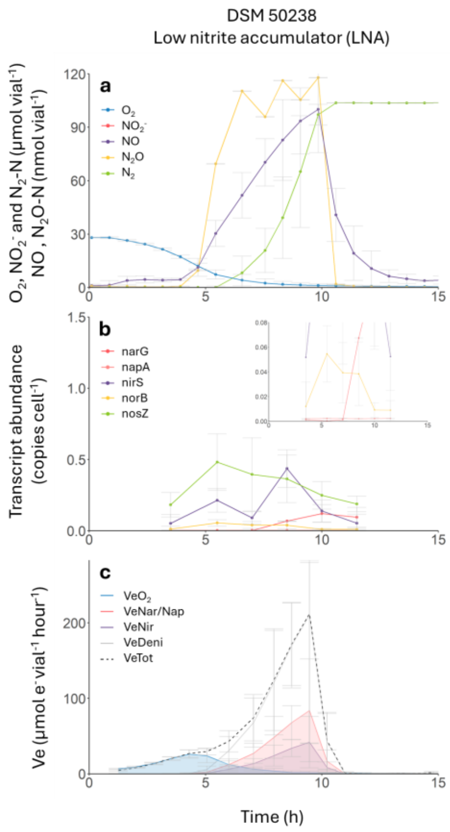


**Figure S6. Denitrification kinetics (a), denitrification gene transcription (b), and electron flow rates to reductases (c) during and after the transition from aerobic respiration to denitrification for the LNA strain DSM 50238.** The strain was incubated at 20 °C in 120 mL medical vials containing 50 mL Sistrom's mineral medium supplemented with $KNO_3^-$ to an initial concentration of 2 mM (100 µmol vial$^{-1}$), with He and 1% $O_2$ in the headspace. Graphs show mean values from three independent replicates (n = 3); error bars indicate standard deviation.

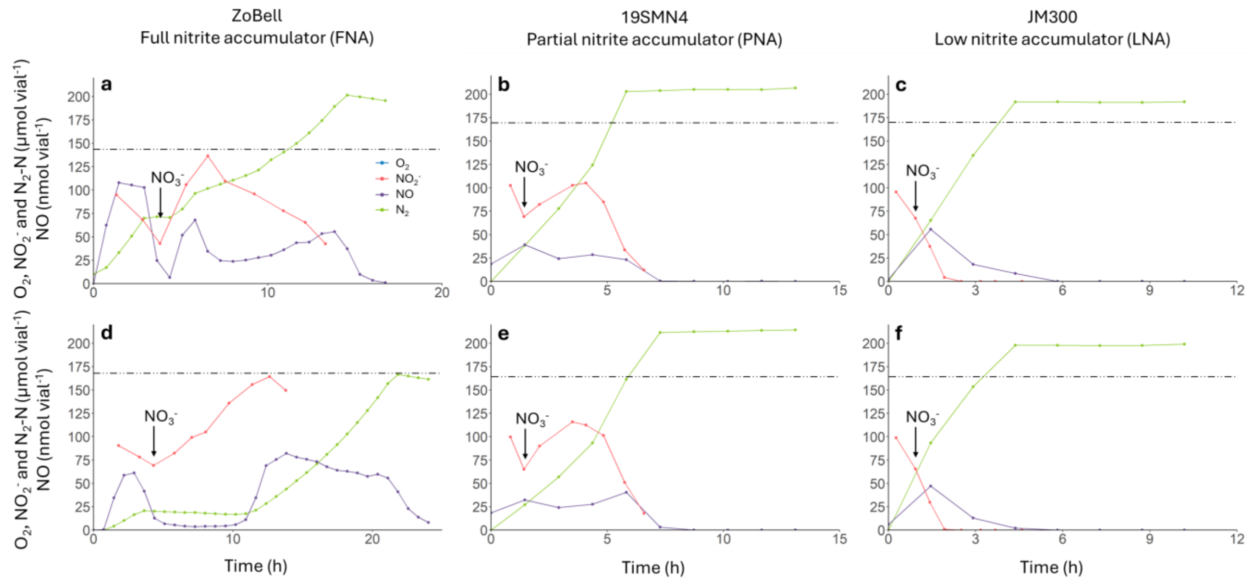


**Figure S7. Competition for electrons between $NO_3^-$ and $NO_2^-$ reduction pathways in three *Stutzerimonas* strains, each representing the three groups of nitrite accumulators**. The timing of $NO_3^-$ addition varied between replicate vials; one representative replicate is shown in Fig. 5a–c of the main text, and the two others are shown here. Cultures were first incubated under anoxic conditions (He headspace) in Sistrom´s medium containing $NO_3^-$, to produce cells with a complete denitrification proteome. At t = 0, cells from these precultures were transferred to fresh anoxic vials containing Sistrom's medium with 2 mM $NO_2^-$ (100 µmol vial$^{-1}$), with no $NO_3^-$ initially present. When approximately one third of the $NO_2^-$ had been reduced, 100 µmol $NO_3^-$ (corresponding to 2 mM in the medium) was added (indicated by the arrow). Dashed lines represent the theoretical maximum accumulation of $NO_2^-$.

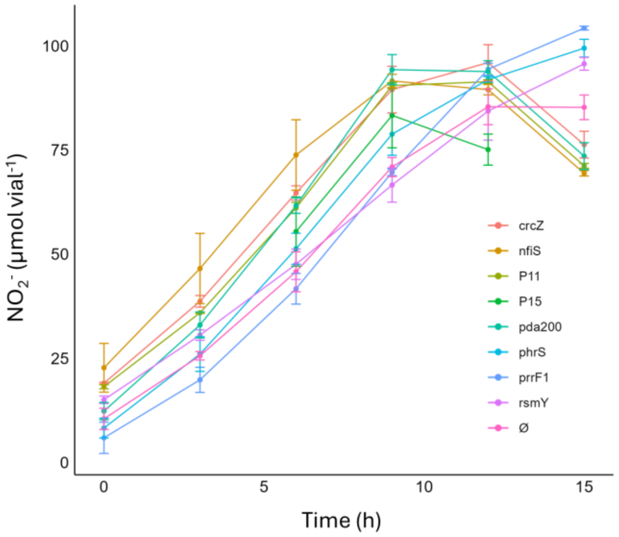


**Figure S8. Anaerobic nitrite accumulation kinetics of *Stutzerimonas perfectomarina* ZoBell sRNA overexpression mutants.** Eight sRNAs (crcZ, nfiS, P11, P15, pda200, phrS, prrF1, rsmY) (Lin et al., 2015; Zhan et al., 2016; Wang et al., 2022) were cloned into the arabinose-inducible vector pL2020 and transformed into *S. perfectomarina* ZoBell. Anaerobic cultures were grown in helium-flushed, sealed vials containing Sistrom's medium supplemented with 30 µg $mL^{-1}$ chloramphenicol and 2 mM nitrate (100 µmol $vial^{-1}$) as the terminal electron acceptor. The cultures were induced by adding arabinose (2 mg $mL^{-1}$) at the beginning of the incubation. Ø represent the *Stutzerimonas perfectomarina* Zobell strain carrying the empty pL2020 vector. Graph shows mean values from three independent replicates (n = 3); error bars indicate standard deviation.

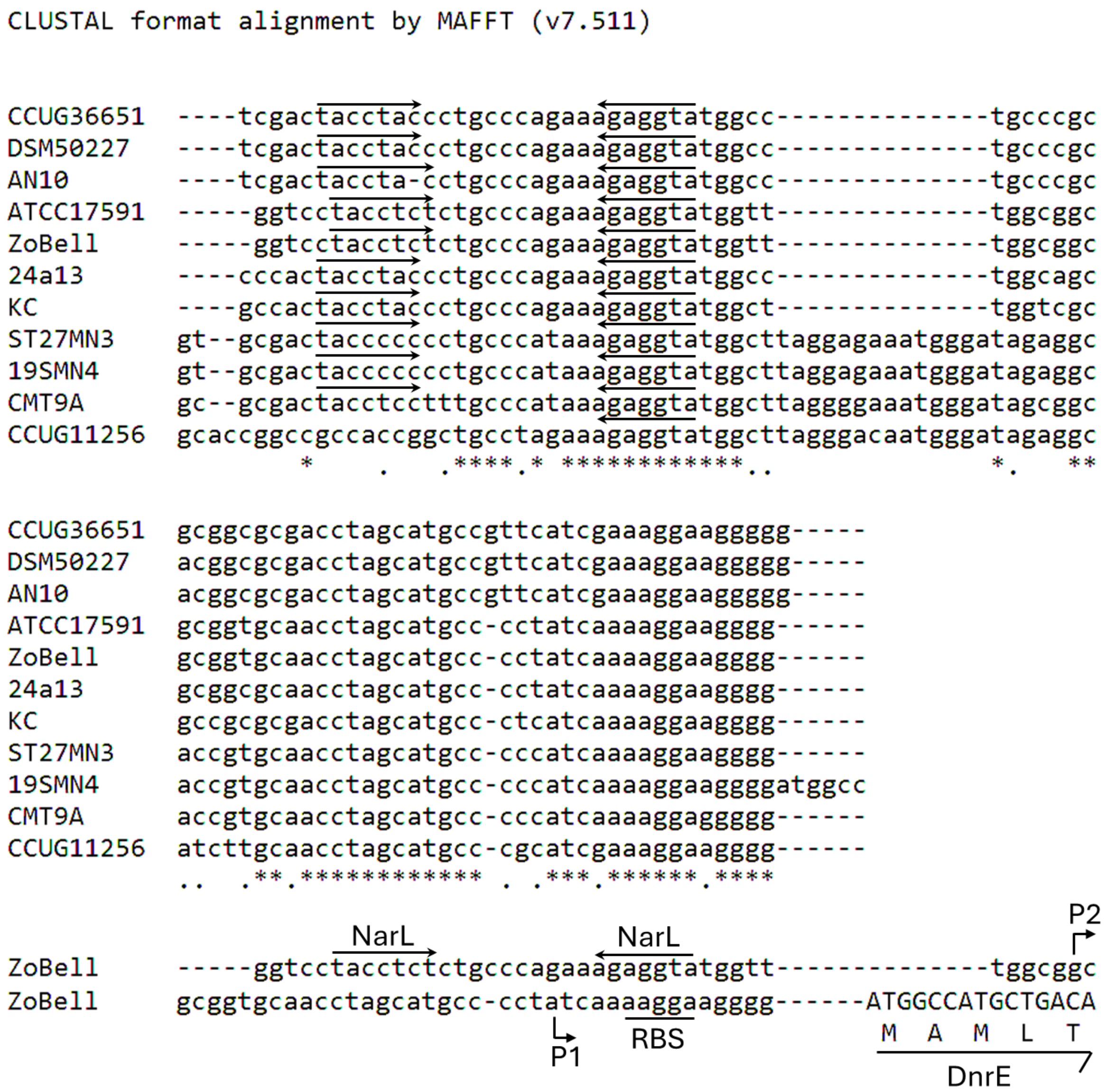


**Figure S9. MAFFT multiple sequence alignment of the intergenic region upstream of *dnrE* in 11 Stutzerimonas strains exhibiting the FNA and PNA phenotypes, all of which carry the dnrE gene.** Potential NarL binding sites are indicated by arrows, with the recognition heptamer consensus sequence in *E. coli* (TACYYMT), where Y represents T or C and M represents C or A (Härtig et al. 1999). Promoters are shown as cornered arrows, with P1 being $NO_3^-$ responsive (Vollack et al. 1999). Ribosome binding sites (RBS) are underlined. The *dnrE* coding sequence is shown in uppercase letters, and the corresponding translated amino acids are indicated above the half arrow.

**Table S1. Description of the *Stutzerimonas* strains used in this study.**

| Group | Strain | Old nomenclature | Genomovar | NCBI RefSeq | Isolation place and environment | References |
|---|---|---|---|---|---|---|
| FNA | *S. chloritidismutans* CCUG 36651 * | *P. stutzeri* CCUG 36651 | 3 | GCF_002890855.1 | Sweden; Water, borehole | Mulet et al., 2008 |
| | *S. chloritidismutans* DSM 50227 * | *P. stutzeri* DSM 50227 | 3 | GCF_002843895.1 | Unknown; Clinical | Van Niel and Allen, 1952 |
| | *S. perfectomarina* ATCC 17591 * | *P. stutzeri* ATCC 17591 | 2 | GCF_002890835.1 | Denmark; Clinical | Stanier et al., 1966 |
| | *S. perfectomarina* Zobell | *P. stutzeri* ZoBell | 2 | GCF_000237885.1 | Pacific Ocean; Marine | ZoBell and Upham, 1944 Peña et al., 2012 |
| | *S. stutzeri* 24a13 * | *P. stutzeri* 24a13 | 16 | GCF_002909485.1 | Germany; Mineral oil-contaminated soil | Sikorski et al., 2002 |
| | *S. stutzeri* KC * | *P. stutzeri* KC | 9 | GCF_002890795.1 | USA California; Aquifer | Sepúlveda-Torres et al., 2001 |
| | *S. decontaminans* ST27MN3 * | *P. stutzeri* ST27MN3 | 4 | GCF_002890955.1 | Spain; Marine | Scotta et al., 2012 |
| | *S. stutzeri* CMT.9.A | *P. stutzeri* CMT.9.A | 1 | GCF_000195105.1 | Germany; Sorghastrum nutans rhizosphere | Krotzky and Werner, 1987 |
| PNA | *S. chloritidismutans* AN10 | *P. stutzeri* AN10 | 3 | GCF_000267545.1 | Spain; Contaminated marine sediment | Bosch et al., 2000 |
| | *S. chloritidismutans* 19SMN4 | *P. stutzeri* 19SMN4 | 4 | GCF_000661915.1 | Spain; Contaminated marine sediment | Rosselló et al., 1991 |
| | *S. stutzeri* CCUG 11256 | *P. stutzeri* CCUG 11256 | 1 | GCF_000219605.1 | Unknown; Spinal fluid | Chen et al., 2011 |
| LNA | *S. frequens* DNSP21 * | *P. stutzeri* DNSP21 | 5 | GCF_002890935.1 | Spain; Wastewater | Rosselló et al., 1991 |
| | *S. stutzeri* 24a75 * | *P. stutzeri* 24a75 | 17 | GCF_002890915.1 | Germany; Mineral oil-contaminated soil | Sikorski et al., 2002 |
| | *S. degradans* | *P. stutzeri* | 7 | GCF_002891015.1 | USA; Soil | Stanier et al., 1966 |

| | | | | | |
|---|---|---|---|---|---|
| DSM 50238 * | DSM 50238 | | | | |
| *S. xanthomarina*<br>DSM 18231 | *P. xanthomarina*<br>DSM18231 | n/a | GCF_900129835.1 | Japan; Marine ascidian | Romanenko et al., 2005 |
| *S. stutzeri*<br>4C29 * | *P. stutzeri*<br>4C29 | 15 | GCF_002890895.1 | Germany; Marine sediment | Sikorski et al., 2002 |
| *S. stutzeri*<br>28a3 * | *P. stutzeri*<br>28a3 | 14 | GCF_002890995.1 | Israel; Soil | Sikorski et al., 2002 |
| *S. nitritolerans*<br>JM300 | *P. stutzeri*<br>JM300 | 8 | GCF_000279165.1 | USA; Soil | Rosselló-Mora et al., 1996<br>Busquets et al., 2012 |

Strains marked with an asterisk (*) were sequenced as part of this study.

**Table S2. Primers used in this study.**

| Primer | Sequence 5'-3' | Description |
|---|---|---|
| Fw *narG* ZoBell | ATCGAGTGCTTCAACGCCAA | Targeting *narG* of the strain ZoBell for qPCR analysis |
| Rv *narG* ZoBell | ATCATGTGGGTGGGCTTGAG | |
| Fw *narG* JM300 | GAAGGGCAAGAAGAGCAACG | Targeting *narG* of the strain JM300 for qPCR analysis |
| Rv *narG* JM300 | CAGGGTGAGCATGATCAGGT | |
| Fw *napA* ZoBell | CGCCAACAAGCTGATCAACA | Targeting *napA* of the strain ZoBell and JM300 for qPCR analysis |
| Fw *napA* JM300 | TTTCTTCGACGCCAACAAGC | |
| Rv *napA* ZoBell and JM300 | TCTTCACGGCGCATTTCTTG | |
| Fw *nirS* ZoBell | CGTGGTATCGCTCATCTCCA | Targeting *nirS* of the strain ZoBell for qPCR analysis |
| Rv *nirS* ZoBell | GGTCTTGACGAACAGGTTGC | |
| Fw *nirS* JM300 | TGGCAGCTCTGATCGATACC | Targeting *nirS* of the strain JM300 for qPCR analysis |
| Rv *nirS* JM300 | TGCTCCTTGTACTTGGCGTA | |
| Fw *norB* ZoBell | CTGTTCGCGTTCTACAACCC | Targeting *norB* of the strain ZoBell for qPCR analysis |
| Rv *norB* ZoBell | GGCGATGATCACGTACAACC | |
| Fw *norB* JM300 | TCCTGATCAAGATCACCGGC | Targeting *norB* of the strain JM300 for qPCR analysis |
| Rv *norB* JM300 | CAGCACCATGGCGAAGAAC | |
| Fw *nosZ* ZoBell and JM300 | CGGTGTTCAACGTCGACTC | Targeting *nosZ* of the strain ZoBell and JM300 for qPCR analysis |
| Rv *nosZ* ZoBell and JM300 | TGTTGGCCTTGTCGTTGATG | |
| Fw *narG* DSM 50238 | TACGTCGGTCAGGAAAAGC | Targeting *narG* of the strain DSM 50238 for qPCR analysis |
| Rv *narG* DSM 50238 | GCGAGCTGTGGTTGTAGAAG | |
| Fw *napA* DSM 50238 | TTCTTCGACGCCAACAAAC | Targeting *napA* of the strain DSM 50238 for qPCR analysis |
| Rv *napA* DSM 50238 | TGCTGACCACTTCGATCTTG | |

| | | |
|---|---|---|
| Fw *nirS* DSM 50238 | TGTGGACAAGCAGGAATACC | Targeting *nirS* of the strain DSM 50238 for qPCR analysis |
| Rv *nirS* DSM 50238 | AGAATCTTGCCGGTTTCCT | |
| Fw *norB* DSM 50238 | TTGACCGTGAAGTGATCGAG | Targeting *norB* of the strain DSM 50238 for qPCR analysis |
| Rv *norB* DSM 50238 | GGTGACCTGTACCGATGATG | |
| Fw *nosZ* DSM 50238 | ACGCCGAGAAGATGGAAAT | Targeting *nosZ* of the strain DSM 50238 for qPCR analysis |
| Rv *nosZ* DSM 50238 | AAGGCCTTCTCCGAGTTGTA | |
| Fw pCasPA | ATTATGTTGGTCCATTGGC | Specific primers of pCasPA |
| Rv pCasPA | CAACCAGTATAACGGCGAC | |
| Fw pACRISPR | ATTAATCATCCGGCTCGTAT | Verify the insertion of sgRNA and repair template |
| Rv pACRISPR | TATGACCATGATTACGCCA | |
| Fw sgRNA *narG* | GTGGGCGGCGCAACATGTCCCCAG | sgRNA sequence for deletion of *narG* |
| Rv sgRNA *narG* | AAACCTGGGGACATGTTGCGCCGC | |
| Fw sgRNA *dnrE* | GTGGACGCAAGGTGTTGAGCACTG | sgRNA sequence for deletion of *dnrE* |
| Rv sgRNA *dnrE* | AAACCAGTGCTCAACACCTTGCGT | |
| Fw up *narG* | TGTCCATACCCATGGTCTAGACCCGGTCGGCACCATCCA | Amplification for 1kb upstream repair of *narG* |
| Rv up *narG* | GGCCTATTCCTGGTTTCTCTCCTCACTCCGGT | |
| Fw down *narG* | AGAGAAACCAGGAATAGGCCATGAAGATTCGTTC | Amplification for 1kb downstream repair of *narG* |
| Rv down *narG* | GGGAGTATGAAAAGTCTCGAGGTAGACCGGCGACTTCTGC | |
| Fw up *dnrE* | TGTCCATACCCATGGTCTAGATGCCGAGGACATGCTCGT | Amplification for 1kb upstream repair of *dnrE* |
| Rv up *dnrE* | AAGGAAGGGGGCCCTACTCGGACGGATCG | |
| Fw down *dnrE* | CGAGTAGGGCCCCCTTCCTTTGATGGGGGC | Amplification for 1kb downstream repair of *dnrE* |
| Rv down *dnrE* | GGGAGTATGAAAAGTCTCGAGACGGCGGTAAGCCCTTGAC | |
| Fw pL2020 PrrF1 | AGGAGGAATTACATAGAGAATTGTTATTATTATCGCAACT | Amplification for the sRNA PrrF1 |
| Rv pL2020 PrrF1 | CAAAACAGCCAAGCTCAGGCCGATTACGTCTGGTACA | |

| | | |
|---|---|---|
| Fw pL2020 PhrS | AGGAGGAATTACATATTGCCGGTTTAACTTGAACCC | Amplification for the sRNA PhrS |
| Rv pL2020 PhrS | CAAAACAGCCAAGCTGATCAGTCGCCTGGCCAC | |
| Fw pL2020 RsmY | AGGAGGAATTACATAACGGTCTGGCGGTAATCTACTG | Amplification for the sRNA RsmY |
| Rv pL2020 RsmY | CAAAACAGCCAAGCTGGCGAAGGTCATTTGCTTCATCG | |
| Fw pL2020 CrcZ | AGGAGGAATTACATAACGACAACTGCTTACTTAATGGC | Amplification for the sRNA CrcZ |
| Rv pL2020 CrcZ | CAAAACAGCCAAGCTGTGTGGCGGAATGACGGG | |
| Fw pL2020 P11 | AGGAGGAATTACATATTCGCACCATGAGGGCGC | Amplification for the sRNA P11 |
| Rv pL2020 P11 | CAAAACAGCCAAGCTATGCCGTTGCCGATCATCACC | |
| Fw pL2020 P15 | AGGAGGAATTACATATCGCCATGCGGGTTGAAA | Amplification for the sRNA P15 |
| Rv pL2020 P15 | CAAAACAGCCAAGCTGGCGTCGAAGCGGCTAAA | |
| Fw pL2020 NfiS | AGGAGGAATTACATAGCGCCGCCAGTCCCACCGAA | Amplification for the sRNA NfiS |
| Rv pL2020 NfiS | CAAAACAGCCAAGCTTCGGGTAGCGCCGCGATTGA | |
| Fw pL2020 pda200 | AGGAGGAATTACATACGCGCCCGGCCTTGATCG | Amplification for the sRNA pda200 |
| Rv pL2020 pda200 | CAAAACAGCCAAGCTTGCAGGCCGAACCGACCTTCATCG | |
| Fw pL2020 paiI | AGGAGGAATTACATACGCGGGACTTAGTCTTGAGC | Amplification for the sRNA paiI |
| Rv pL2020 paiI | CAAAACAGCCAAGCTCAGGCACCCTTCGGTACTCG | |

**Table S3. Bacterial strains and plasmids used for generating *Stutzerimonas decontaminans* deletion mutant and the *Stutzerimonas perfectomarina* overexpression mutants in this study.**

| Strain | Description | Source |
|---|---|---|
| *E. coli* DH5α | Cloning strain | Lab stock |
| *S. decontaminans* 19SMN4 | Wild-type | DSM |
| 19SMN4 pCasPA | 19SMN4 harboring pCasPA | This study |
| 19SMN4 *ΔnarG* | 19SMN4 *narG* gene deleted | This study |
| 19SMN4 *ΔdnrE* | 19SMN4 *dnrE* gene deleted | This study |
| *S. perfectomarina* ZoBell | Wild-type | ATCC |
| ZoBell PrrF1 | ZoBell harboring pL2020 PrrF1 | This study |
| ZoBell PhrS | ZoBell harboring pL2020 PhrS | This study |
| ZoBell RsmY | ZoBell harboring pL2020 RsmY | This study |
| ZoBell CrcZ | ZoBell harboring pL2020 CrcZ | This study |
| ZoBell P11 | ZoBell harboring pL2020 P11 | This study |
| ZoBell P15 | ZoBell harboring pL2020 P15 | This study |
| ZoBell NfiS | ZoBell harboring pL2020 NfiS | This study |
| ZoBell pda200 | ZoBell harboring pL2020 pda200 | This study |
| ZoBell paiI | ZoBell harboring pL2020 paiI | This study |
| Plasmids | Description | Reference |
| pCasPA | Tetracyline resistant, for expression of Cas9 protein and λ-Red recombination system | (Chen et al., 2018) |
| pACRISPR | Carbenicillin resistant, for expression of sgRNA and assembling homologous repair arms | (Chen et al., 2018) |
| pACRISPR *ΔnarG* | pACRISPR derivative for *narG* deletion | This study |
| pACRISPR *ΔdnrE* | pACRISPR derivative for *dnrE* deletion | This study |
| pL2020 | Chloramphenicol resistant, araBAD based expression vector for inducible protein production | (Sommer et al.,2017) |
| pL2020 PrrF1 | pL2020 derivative carrying the sRNA PrrF1 | This study |
| pL2020 PhrS | pL2020 derivative carrying the sRNA PhrS | This study |
| pL2020 RsmY | pL2020 derivative carrying the sRNA RsmY | This study |
| pL2020 CrcZ | pL2020 derivative carrying the sRNA CrcZ | This study |
| pL2020 P11 | pL2020 derivative carrying the sRNA P11 | This study |
| pL2020 P15 | pL2020 derivative carrying the sRNA P15 | This study |
| pL2020 NfiS | pL2020 derivative carrying the sRNA NfiS | This study |
| pL2020 pda200 | pL2020 derivative carrying the sRNA pda200 | This study |
| pL2020 paiI | pL2020 derivative carrying the sRNA paiI | This study |

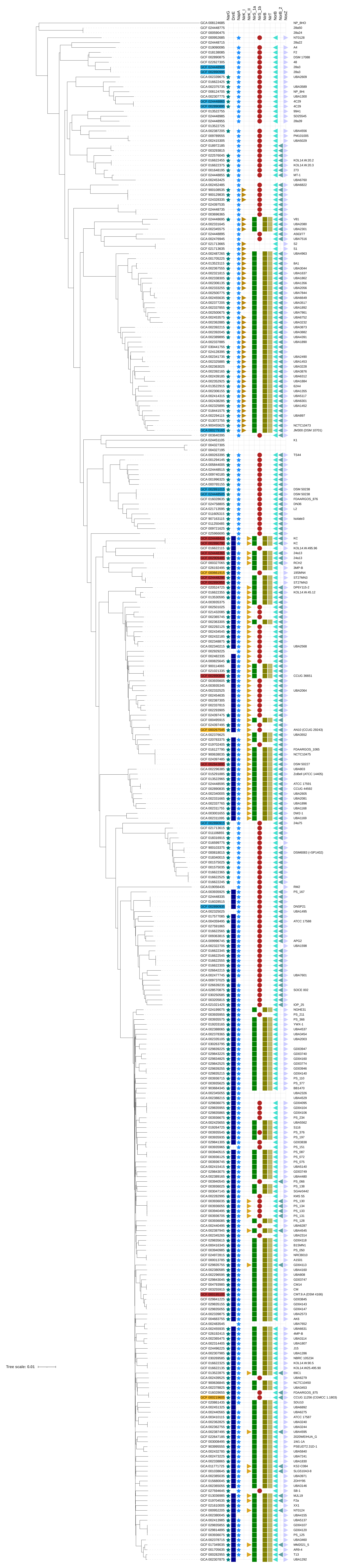
NarG
DnrE
NapA
NirK_I
NirK_II
NirS_1a
NirS_1b
NirB
NirT
NorB
NorB_2
NosZ
Tree scale: 0.01

**Figure S4.** ***Stutzerimonas*** **genome clustering tree and $NO_2^-$ accumulation phenotypes of the strains examined in this study**. The three $NO_2^-$ accumulation groups are indicated by different colors in the phylogenetic tree. Red: Full $NO_2^-$ accumulators (FNA); Yellow: Partial $NO_2^-$ accumulators (PNA); Blue: Low $NO_2^-$ accumulators (LNA). Selected genes associated with denitrification are indicated. To generate the tree an extensive collection of genomes from organisms given the genus name *Stutzerimonas* was gathered. This was done using the advanced search in GTDB (https://gtdb.ecogenomic.org/). The search term was g__Stutzerimonas and the filters used were CheckM completeness > 95%, CheckM contamination < 10%, and number of contigs < 80. At the time this was run (October, 2024) this returned 322 genomes. After manual filtering to remove small genomes and genomes with other issues a final set of 314 genomes was downloaded from NCBI using the curl file generated by GTDB. Some strains have been sequenced multiple times, but these were not trimmed to a single genome, therefore, some strains appear more than once in the output. These genomes were clustered based on k-mer distances using PopPUNK 2.6.0 (POPulation Partitioning Using Nucleotide Kmers) (Lees et al., 2019). For creating the database these command line options were used: --length-sigma 1 --plot-fit 3 --min-k 17 --max-k 41 --sketch-size 100000. Cluster fitting was performed using dbscan with a --K of 6 and the output from that then refined. The output of this refinement was used to produce a neighbor-joining tree, which was uploaded to the Interactive Tree of Life (https://itol.embl.de/) (Letunic and Bork, 2024) and modified there.
